# Neural Substrates of Approach–Avoidance Control in Motivational Conflict

**DOI:** 10.1101/2025.10.30.685670

**Authors:** Menghuan Chen, Janna Teigeler, Paul Pauli, Andre Pittig, Matthias Gamer

## Abstract

Adaptive behavior requires control over automatic tendencies to approach reward and avoid threat. Dysregulation of this control characterizes disorders such as anxiety or addiction, yet the underlying neural mechanisms remain unclear. Here, we investigated circuits engaged during approach-avoidance control using fMRI with concurrent eye-tracking, and behavioral measures. Forty participants (22 females) completed an approach-avoidance conflict task with free- and forced-choice trials. Each trial comprised anticipation, response (using a joystick to approach/obtain or avoid/forgo outcomes), and outcome phases. Before scanning, participants learned that conditioned stimuli (CS) predicted an aversive stimulation (avCS+), appetitive reward (appCS+), both outcomes (confCS+), or no outcome (neuCS−). Discordant responses (e.g., approaching avCS+) were slower than concordant responses (e.g., avoiding avCS+), confirming heightened control demands. Overcoming threat-driven avoidance specifically recruited the left inferior frontal gyrus (IFG), while suppressing reward-driven approach lacked distinct neural signatures. During anticipation, confCS+ and avCS+ showed overlapping activation in salience–control networks, including middle/anterior cingulate cortex (MCC/ ACC), anterior insula, IFG, and ventral striatum (VS). Interestingly, confCS+ evoked threat-like anticipatory neurophysiological responses (pupil dilation; ACC/insula activation) but subsequently triggered reward-like approach behavior. Multivariate pattern and psychophysiological interaction analyses revealed that this dissociation was driven by differential encoding of upcoming responses in the ventral striatum during the anticipation phase and by altered functional coupling between the VS and the right temporoparietal junction (rTPJ). These findings indicate that motivational control prioritizes salience over valence and suggest a VS-rTPJ network for resolving approach-avoidance conflicts, offering novel insights into neural dynamics of flexible goal-directed behavior.

**Significance Statement:** This study advances our understanding of how the brain dynamically resolves conflicts between approaching rewards and avoiding threats. By measuring neural activity during approach-avoidance conflicts, our study identified overlapping neural circuits for threat and conflict processing (e.g., middle/anterior cingulate, ventral striatum). It revealed that threat-like neural responses to conflicting stimuli triggered approach behaviors resembling reward-motivated actions. Critically, connectivity between the ventral striatum and the temporoparietal junction predicted individual differences in conflict resolution, linking network-level interactions to adaptive behavioral control. These findings suggest salience overrides valence in motivational control and motivate hypothesis-driven tests of network disruption in psychiatric populations.

## Introduction

Automatic tendencies to approach rewards and avoid threats are critical for survival, as they enable rapid responses in dynamic environments. Yet, adaptive behavior often requires overcoming these reactions, such as approaching feared stimuli (e.g., medical procedures) or avoiding desired ones (e.g., addictive substances) to align actions with long-term goals, particularly under conflict where competing motivations arise (Shenhav et al., 2013; Soutschek et al., 2014). Dysregulated control over such approach-avoidance biases has been implicated in various psychopathologies, e.g., anxiety and addiction (Loijen et al., 2020). Behavioral interventions commonly aim to retrain individuals to overcome these automatic responses (Kakoschke et al., 2017; Pittig et al., 2021). However, the neural mechanisms that support such control in the face of motivational conflict remain incompletely understood.

Human neuroimaging studies have revealed that affective valuation systems bias behavior through distinct but interacting neural pathways. The amygdala amplifies aversive predictions to promote avoidance (Davis et al., 2010; Wen et al., 2022), whereas the ventral striatum (VS) encodes expected reward value to facilitate approach (Goldstein et al., 2012; Roesch et al., 2009). However, functional overlap is observed as both areas also contribute to opposite responses (Murty et al., 2023; Wassum, 2022). Connectivity studies show that amygdala– dorsomedial prefrontal coupling predicts avoidance under threat, and striatal–prefrontal coupling during reward tracks approach, underscoring the integration of motivational signals with top-down control (Dobryakova & Smith, 2022; Gold et al., 2015; Leitão et al., 2022). These processes are accompanied by robust autonomic signatures, as both reward and threat anticipation reliably elicit sympathetic arousal (Merscher et al., 2022; Schneider et al., 2018). Under reward-threat conflict, behavior emerges from interactions among valuation hubs and salience/control networks. The salience network, anchored by the anterior cingulate cortex (ACC) and anterior insula, detects motivational conflict and salience, arbitrates between competing actions, and engages control circuits (Botvinick et al., 2001; Seeley et al., 2007; Shenhav et al., 2013). Recent work demonstrates that subcortical nodes (e.g., VS, thalamus) dynamically integrate reward and threat information prior to approach-avoidance decisions (Hulsman et al., 2024). The middle cingulate cortex (MCC) associates actions with resulting outcomes, projecting to premotor regions to support action selection and initiation (Procyk et al., 2016). The prefrontal cortex (PFC) underpins cognitive control and is modulated by affective states (Bari & Robbins, 2013; Grahek et al., 2020; Levy & Wagner, 2011). Classic tasks (e.g., Go/No-Go task) implicate the inferior frontal gyrus (IFG) in suppressing prepotent responses to enable goal-directed action (Guitart-Masip et al., 2012), while emerging evidence highlights the anterior PFC (aPFC) in emotional control, with perturbation impairing the capacity to overcome affective impulses (Bramson et al., 2018; Volman et al., 2011). Despite these insights, it remains unclear how motivational salience is integrated with cognitive control to guide approach-avoidance decisions.

The current study sought to elucidate the neural and cognitive processes underlying control over automatic approach-avoidance behaviors. To this end, we combined fMRI, eye-tracking, and behavioral measures with a previously established approach-avoidance conflict task, in which conditioned stimuli predicting appetitive, aversive, conflicting, and neutral outcomes (appCS+, avCS+, confCS+, and neuCS−) were presented in an anticipation phase followed by free versus forced approach-avoidance responses (Chen et al., 2025). Based on the role of the salience network in prioritizing motivationally relevant stimuli, we hypothesized that affective stimuli would evoke stronger activation of the salience network (ACC, anterior insula) than the neutral condition during anticipation, and that the confCS+ would elicit stronger salience engagement relative to pure reward or threat cues, respectively, reflecting its heightened motivational ambiguity and demand for conflict processing. Discordant responses requiring suppression of prepotent tendencies (i.e., approaching avCS+, and avoiding appCS+) were expected to engage control networks (IFG and aPFC) more strongly than concordant responses. Finally, we expected anticipatory connectivity between affective valuation hubs (e.g., ACC, VS) and control regions to be stronger during conflict than for pure threat/reward anticipation, and this coupling would correlate with behavioral choices (e.g., approaching confCS+ despite concurrent threat signals).

## Materials and Methods

### Participants

Sample size planning was informed by the behavioral effects of our previous study employing the same experimental paradigm (Chen et al., 2025). The present study relies on a 4 × 2 within-subjects design (4 CS types × 2 response types) and was specifically powered to detect an interaction between these factors using a simulation-based approach (Lakens & Caldwell, 2021). With a targeted sample size of *N* = 40 participants, statistical power exceeded 0.95 at an α-level of 0.05. In total, 48 participants were recruited to account for dropouts. Of these, one participant withdrew during the imaging assessment due to excessive distress. Seven of the remaining 47 participants who completed the AAC task during fMRI were excluded from analyses because of scanner technical issues (*n* = 5) or procedural errors (*n* = 2). This yielded a final sample of 40 participants (22 females, mean age ± *SD* = 25.15 ± 4.94 years, range 19–39 years). The demographic and questionnaire data are provided in Table S1. Inclusion criteria were an age range of 18–40 years, right-handedness, and normal or corrected-to-normal vision with contact lenses. All participants were screened to be free from self-reported current or lifetime diagnosis of any psychiatric disorder, substance abuse, major medical and neurological disorders, pain-related diseases, or MR-safety risks. Participants received 13 € per hour for participation and an additional bonus of up to 5 € depending on their performance in the task. All participants provided written informed consent prior to completing the study. All procedures were approved by the Institutional Review Board at the University of Würzburg (GZEK 2023-53). Raw behavioral and eye-tracking data, participant-level GLM beta images, and supplementary information are shared via the Open Science Framework (https://osf.io/j5hze).

### Procedure

After providing written informed consent, participants first completed several questionnaires on individual differences (see Table S1) that might influence task performance outside the scanner. Sociodemographic data included age, sex, education, and health-related behaviors (e.g., cigarette, coffee, and alcohol consumption and physical activity). Questionnaires assessed intolerance of uncertainty (IU-18) (Dugas et al., 2004; Gerlach et al., 2008), depression, anxiety and stress (German version of DASS-21) (Lovibond & Lovibond, 1995; Nilges & Essau, 2015), behavioral inhibition and approach system (BIS/BAS) (Carver & White, 1994), personality (Rammstedt & John, 2007), and general decision-making style (Scott & Bruce, 1995).

Afterwards, participants were positioned in the scanner in a supine position. Two stainless-steel surface electrodes (9-mm diameter; GVB-geliMED, Bad Segeberg, Germany) for delivering aversive electrical stimulation were attached to the inner side of the participant’s left calf, and the joystick mounted on a wooden stand was placed next to the right upper leg. Participants were asked to test the joystick’s movement and adjust the stand’s position to ensure it was the most comfortable for them to respond. Next, participants underwent a calibration procedure to determine their pain threshold intensity for the electrical stimulation (aversive unconditioned stimulus, avUS). The stimulation consisted of 3 electrical pulses (2-ms pulse duration separated by 48 ms) and was generated by a constant current stimulator (Digitimer DS7A, Digitimer Ltd., Welwyn Garden City, UK). The intensity was calibrated for each participant, using a staircase procedure consisting of two ascending and descending series of stimulation to achieve a perceived avUS rating of 6 (*clearly painful but tolerable*) on a scale from 0 (*not painful at all*) to 10 (*very painful*). The procedure used has been described in previous studies (Andreatta et al., 2010; Reutter & Gamer, 2023). Calibration resulted in a mean final intensity of 9.19 mA (*SD* = 8.21 mA).

A reward calibration procedure for each participant was conducted immediately after the avUS calibration to determine the amount of competing reward (appetitive unconditioned stimulus, appUS). During this procedure, participants were presented with a series of questions “Are you willing to pay_cents in order to avoid the electrical stimulation?”. The reward amounts ranged from 4 to 30 cents in even increments (i.e., 4, 6, …, 28, 30 cents) and were presented in a randomized order. Participants were required to answer either “yes” or “no” to each question. The competing reward amount was calculated as the average of the lowest amount that received a “no” and the highest amount that received a “yes”. Similar procedures were used in our previous studies (Chen et al., 2025; Wong & Pittig, 2022). This resulted in a mean competing reward of 9.83 cents (*SD* = 8.77 cents). The competing reward was designed to balance the incentives of reward-seeking and threat-avoidance in trials with conflicting reinforcement (confUS). Participants were instructed that the money gained in each trial was aggregated (up to a maximum of 5 €) and paid at the end of the experiment.

Participants then accomplished a short acquisition training with two trials per condition to acquire the associations between conditioned stimuli (CSs) and unconditioned stimuli (USs). Since this study did not aim to investigate contingency learning, participants were explicitly informed of the relationship between CSs and USs and were asked to remember the relationship as accurately as possible. Four neutral geometric shapes of equal size and brightness served as 4 CS types. During the acquisition training, one shape served as the aversive CS+ (avCS+) and was always followed by the aversive US. A second shape was the appetitive CS+ (appCS+) and always followed by the appetitive US. A third shape served as the conflicting CS+ (confCS+) and was concurrently followed by both the aversive and appetitive US. Finally, the fourth shape served as the neutral CS (neuCS−) and was never followed by any USs. The association between a specific CS and the different outcomes was counterbalanced across participants. CSs were presented for 6 s in a randomized order, and the corresponding US was delivered 4 s after CS onset, with an inter-trial-interval (ITI) varying randomly between 3, 4, or 5 s.

Next, participants completed six practice trials to familiarize themselves with the joystick movement in the scanner and to ensure full understanding of the task. Participants then completed the free versus forced Approach-Avoidance Conflict (AAC) task (see below and Figure 1) in the MRI scanner, which was divided into three blocks. Before the first block, participants received explicit instructions about the association between each CS and the corresponding US. To ensure that the electrical stimulation was sufficiently intense and to recalibrate it, if necessary (i.e., when the rating was ≤ 5), participants were asked to rate the pain intensity of the electrical stimulation after each block of the AAC task. Across the imaging session, 90% of participants underwent at least one intensity adjustment to sustain the target aversive rating. After completing the AAC task, participants rated the valence and arousal of the stimuli outside the scanner on a visual analog scale from 0 (*very unpleasant/calm*) to 10 (*very pleasant/exciting*) and evaluated the corresponding CS-US contingencies from 0 (*very unlikely*) to 100 (*very certain*). They also completed a questionnaire that involved rating on a 0–10 Likert scale, (1) how difficult it was for them to make decisions during the task, (2) how motivated they were to obtain the monetary reward, and (3) how motivated they were to avoid the aversive electrical stimulation.

**Figure 1.**
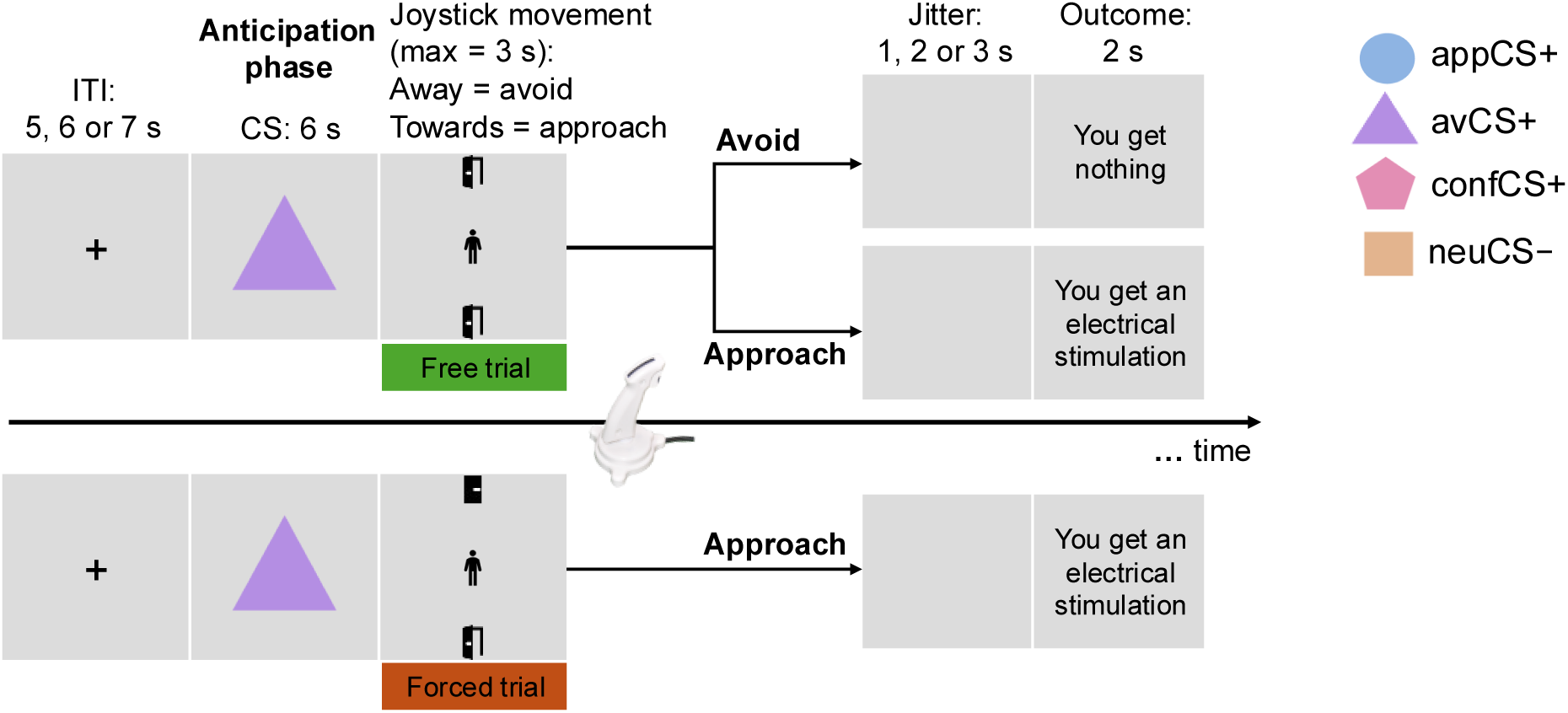
Timeline of an exemplary trial in the free versus forced Approach-Avoidance Conflict (AAC) Task. In each trial, participants were first presented with a fixation cross for 5, 6, or 7 s. Next, a geometric shape (i.e., the CS) was displayed for 6 s during the anticipation phase, where participants freely viewed the stimulus. Following this, a manikin appeared in the center of the screen along with open and/or closed doors at the top and bottom, where participants had up to 3 s to select their choice by moving the manikin into an open door using a joystick. In free trials, both doors were open, and participants were free to decide whether to approach or avoid by pulling the joystick towards themselves (the manikin would be moved into the bottom-open door) or pushing it away (the manikin would be moved into the top-open door), respectively. In forced trials, however, only one door was open, so participants were forced to approach (i.e., only the bottom door was open) or avoid (i.e., only the top door was open). Approach responses led to an outcome determined by the CS type presented during anticipation, while avoidance responses resulted in no outcome, regardless of the CS type. After the manikin reached the open-door area, a jittered blank screen (1–3 s) preceded the outcome display that presented the trial outcome as text for 2 s. The electrical stimulation was immediately delivered after the blank screen for shock trials. Importantly, participants were unaware whether the trial was free or forced until the response phase began. All free and forced trials were mixed and presented in a random order. Please note that the legend in the upper right illustrates one example of the mapping between shapes and CS types; these assignments were counterbalanced across participants.

### Approach-Avoidance Conflict task

During fMRI scanning, participants performed the Approach-Avoidance Conflict (AAC) task (Figure 1), alternating free trials (to probe natural behavioral tendencies) and forced trials (to isolate inhibitory control demands) across four stimulus types. The AAC task was programmed using Presentation Software (Neurobehavioral Systems Inc., Berkeley, California, USA), which was adapted from our previous study (Chen et al., 2025).

Specifically, each trial consisted of three phases. In the anticipation phase, a stimulus (CS) was presented in the center of the screen for 6 s. Four CS types signaled the respective outcomes of monetary reward, aversive stimulation, conflicting outcomes with both the reward and stimulation, and nothing. Participants were instructed to view the stimulus freely during the anticipation. This was followed by the response phase, where a manikin appeared in the center of the screen with an open and/or closed door at the top and the bottom. Participants responded by moving a joystick with their right hand. Notably, the doors were either displayed as “open” or “closed”, which was used to establish free versus forced trials. In free trials, two open doors were displayed, indicating that both approach and avoidance responses were available. Participants could freely choose whether to approach and obtain the CS-specific outcome by pulling the joystick towards them to move the manikin into the bottom-open door or to avoid without receiving anything by pushing the joystick away to move the manikin into the top-open door. In forced trials, only one option was available (i.e., only one door was open), thus participants were forced to perform the only available response. If they tried to perform the unavailable response (i.e., movement towards a closed door), the manikin would not move. Participants had a maximum of 3 s to complete the movement. Once the manikin reached the available open-door area, a blank screen was shown for a random duration of 1 – 3 s. Afterward, the outcome was presented for 2 s, followed by an ITI varying randomly between 5, 6, or 7 s. If participants did not respond within 3 s in particular trials, they received the feedback “You missed the trial”. Participants were instructed to respond as fast as possible. To motivate participants not to miss trials, participants were explicitly informed that missing trials would reduce the total bonus they could earn. Importantly, during the anticipation phase, participants had no information about whether the trial was free or forced, which was only disclosed when the response phase started.

The AAC task included 120 trials, which were presented in a pseudorandomized order for each participant, with no more than three consecutive trials featuring the same CS type or response type. The association between CSs and USs was counterbalanced across participants. Each CS was presented in 30 trials, 10 trials of which were free trials. Among the 20 forced trials, 10 were forced approach, and 10 were forced avoidance. At the beginning of each trial, the joystick’s position was initialized to its neutral position at (0, 0). The AAC task was divided into three blocks of 12 minutes each, with at least one minute of rest in between.

All stimuli were presented on a grey background (RGB: 210, 210, 210) in a central position on a monitor (32-inch full color MR Safe 32 LCD Display with a resolution of 1920 × 1080 pixels and a refresh rate of 120 Hz; BOLD screen 32, Cambridge Research Systems Ltd., UK) located behind the scanner. Participants could see the monitor through a mirror attached to the MRI head coil. Responses were executed with an MR-compatible joystick (Tethyx, Current Designs Inc., Philadelphia, USA) mounted on a wooden stand that allowed for continuous movement of the manikin. Movement trajectories were sampled at 60 Hz during the approach-avoidance conflict task.

During the task, eye movements and pupil diameter were continuously recorded from the participants’ right eye using an MR-compatible EyeLink 1000 Plus system (SR Research Ltd., Ottawa, Canada) with a sampling rate of 250 Hz. Camera and infrared illumination were positioned below the presentation monitor at a fixed distance of 102 cm between the camera and the eye. The eye-tracker was calibrated and validated using a 9-point grid before each block of the task (5-point calibration and validation were used for *n* = 3 participants due to the difficulty in pupil recognition). Out of the 40 participants with valid fMRI imaging data, eye-tracking data of 3 participants were not successfully recorded due to technical difficulties in detecting the pupil.

### Behavioral data analyses

In the AAC task, the proportion of avoidance (P(avoid)) in free trials and response times (RTs) in forced trials were calculated as dependent variables. RTs were measured as the duration between the onset of the manikin and its arrival at the available door area. The continuous movement trajectories of the joystick were also recorded. Omitted trials, in which participants failed to respond within 3 s, were excluded from all further analyses (1.5%; more details about omissions are provided in Supplementary Figure S2). Trials with a joystick y-position outside the neutral range (±10 pixels from [0, 0]) at manikin onset, indicating participants held the joystick in a biased position before the response phase, were also excluded from all further analyses (0.6%). We also assessed whether measures based on the continuous joystick movements, including response latency, execution time, and trajectory length, differed as a function of the experimental manipulations and the participants’ behavior (see Figure S2 for results).

For P(avoid), we conducted a one-way analysis of variance (ANOVA) with the within-subject factor CS type (appCS+, avCS+, confCS+, and neuCS−). Consistent with our previous study (Chen et al., 2025), we expected very few discordant responses (e.g., approaching avCS+) in free trials. Therefore, free trials would not provide sufficient variance to compare approach versus avoidance RTs across all different CSs. Thus, RTs were only compared during forced trials using a 4×2 rmANOVA with CS and response types (forced approach and forced avoidance) as within-subject factors. Greenhouse-Geisser corrections were applied when the sphericity assumption was violated. Significant effects and interactions were followed up with simple main effects and Bonferroni-corrected pairwise *t*-tests. Effect size estimates are reported as partial η^2^ (η_p_^2^) for rmANOVAs and Cohen’s *d* for *t*-tests. JASP software (version 0.19.3.0) was used for statistical analyses.

### Eye-tracking data analyses

The recorded eye-tracking data were converted into MAT files using Edf2Mat Matlab Toolbox (Etter & Biedermann, 2016) in the MATLAB environment (version R2022b; MathWorks, Inc., Natick, Massachusetts, USA). Gaze data were then parsed into saccades and fixations. Saccades were defined as fast eye movements with a velocity exceeding 30° s^-1^ or an acceleration exceeding 8000° s^-2^. Periods between saccades were defined as fixations.

### Pupil diameter

In order to exclude participants with suboptimal data quality, we assessed the number of blink- or eye-closure-related missing pupil values in the raw data. Time series of pupil data were segmented into trials, each consisting of a 1-second baseline before CS onset and the 6-second CS presentation. Trials containing more than 50% of missing data were marked as invalid and excluded from all further analyses (*M* = 2.8%, *SD* = 4.9%). Furthermore, participants with more than half of the total trials marked as invalid were excluded from data analyses (*n* = 1), leaving 36 participants for analyses involving eye-tracking data. Subsequently, we converted the pupil diameter from arbitrary units to millimeter (Hayes & Petrov, 2016). Missing values of the resulting pupil signal were linearly interpolated on a trial-by-trial basis, and the entire tracing was smoothed using a zero-phase low-pass filter with a cutoff frequency of 4 Hz (Kret & Sjak-Shie, 2019). After this procedure, the data were visually inspected to ensure all artifacts were successfully removed. We then calculated changes in pupil diameter relative to the baseline period. The resulting values were then averaged into 12 bins of 0.5 s each, spanning the CS duration.

### Vertical gaze position

Consistent with our previous study (Chen et al., 2025), vertical (but not horizontal) screen scanning was mainly needed to detect the availability of response options in the AAC task. Therefore, here we only analyzed the vertical gaze position to assess visual exploration during the anticipation phase. Information about the horizontal gaze position and fixations is provided in Supplementary Figure S3 and Table S6.

The last 300 ms prior to CS onset were considered as baseline and used for drift correction. To ensure that participants fixated on the center of the screen at trial onset, an iterative drift correction algorithm was applied to the baseline gaze data separately for x and y coordinates. Specifically, the highest and lowest values of baseline coordinates were temporarily removed from the distribution of the baseline data and were marked as invalid when they deviated more than three standard deviations from the mean of the remaining baseline data. This procedure was repeated until no further x- and y-values were marked as invalid. For trials with invalid baseline fixation data, the baseline was replaced using the mean x- and y-coordinates of the remaining trials with valid baseline fixation data (14.04%). Individual gaze drift was finally corrected by subtracting the x- and y-coordinates of the baseline, respectively, from the gaze coordinates during CS presentation for each trial. Similar procedure was established in previous studies (Merscher et al., 2022; Rösler & Gamer, 2019). Finally, relative vertical gaze position was averaged on a second-by-second basis.

To examine condition-related differences in pupil and fixation dynamics during the anticipation phase, we conducted repeated-measures ANOVAs focusing on the final 3 seconds of the CS presentation window (i.e., 3–6 s post-CS onset). By restricting the analyses to the latter portion of the anticipation period, we aimed to isolate sustained task-related modulation linked to the experimental manipulations. Accordingly, 4 × 6 and 4 × 3 rmANOVAs were conducted for pupillary and oculomotor responses, respectively, with CS type and time (the last 3 seconds of CS presentation, divided into 6- or 3-time bins) as within-subject factors. To specifically compare each affective condition (i.e., appCS+, avCS+, confCS+) to the neutral condition (neuCS−), post hoc *t*-tests were performed using false discovery rate correction (Benjamini & Hochberg, 1995) to adjust for alpha-error accumulation.

### fMRI acquisition and analyses

MRI data were acquired using a 3T Skyra MR scanner system (Siemens Medical Systems, Erlangen, Germany) with a 32-channel head coil. Foam inserts were used to fixate the participant’s head within the head coil and to mitigate potential motion artifacts. The complete scan protocol was about 50 min long, including a localizer scan, a field map scan, three functional T2* runs, and a structural T1 scan. After the localizer run, the geometric distortion sensitive field map volumes were acquired (TR = 428 ms; TE 1 = 4.92 ms; distance factor = 30%; number of slices = 42; slice thickness = 2.3 mm; base resolution = 64; field of view (FoV) = 220 mm; flip angle = 60°; voxel size = 3.4×3.4×2.3 mm). For the three functional runs, a multiband sequence was used to collect T2*-weighted echoplanar imaging (EPI) volumes (multiband acceleration factor = 2; TR = 1340 ms; TE = 25 ms; distance factor = 50%; number of slices = 42; slice thickness = 2 mm; flip angle = 60°; in-plane resolution = 2×2 mm; FoV = 216×216 mm^2^; acquisition matrix = 108×108, voxel size = 2×2×2 mm). Finally, T1-weighted structural scans were acquired (TR = 2300 ms; TE = 2.96 ms, distance factor = 50%; slice thickness = 1 mm; base resolution = 256, FoV = 256×256 mm^2^, flip angle = 9°, voxel size =1×1×1 mm).

The fMRI data were processed following the standard preprocessing procedures and analyzed using SPM12 (www.fil.ion.ucl.ac.uk/spm) implemented in MATLAB. After excluding the first five EPI scans of each run to reach steady magnetization, the functional images were slice-timing corrected (temporally aligned to the middle image), realigned and unwarped based on individual voxel displacement maps derived from field mapping to reduce signal drop-out and spatial distortion due to magnetic field inhomogeneities. Following co-registration of the structural and functional scans, the structural scan was segmented and resulting normalization parameters were used to normalize functional volumes to the standard Montreal Neurological Institute (MNI) space. During this step, images were resliced to a voxel size of 2 × 2 × 2 mm, and spatial smoothing with a Gaussian kernel of 6 mm full width at half maximum was applied. Temporal preprocessing involved high-pass filtering at 1/128 Hz and applying a first-order autoregressive model as implemented in SPM12 as default.

#### First level analyses

We constructed event-related general linear models (GLMs) to examine the distinct and shared brain regions related to reward, threat, and conflict anticipation, as well as brain regions underlying overcoming behavioral tendencies (i.e., in trials where participants were forced to approach the threat or avoid the reward). For each trial, we fitted one regressor for the anticipation phase (i.e., 6 s for CS presentation), one regressor for the response phase (i.e., RT, from manikin onset until its arrival at the available door area), and one regressor for the outcome phase separately for the different experimental conditions. This resulted in 4 regressors for the anticipation phase (i.e., 4 CS types, appCS+, avCS+, confCS+, and neuCS−), a maximum of 16 regressors for the response phase (4 CS types × 4 response types, i.e., free approach, free avoidance, forced approach, and forced avoidance). To avoid overparameterization, response regressors were excluded for combinations lacking behavioral instances (e.g., if one participant never freely approached the electrical stimulation, its corresponding regressor was omitted). Eight regressors were included for the outcome phase (i.e., getting the reward, the electrical stimulation, and the conflicting outcomes, respectively, and no outcome due to approach to neuCS−, and no outcome due to avoidance to appCS+, avCS+, confCS+, and neuCS−, respectively). To account for motion-related signal changes, 12 nuisance regressors related to head motion were included: z scores of the three regressors related to translation and three regressors related to rotation of the head, as well as their derivatives. These mentioned regressors were concatenated for the three scanning runs. Lastly, three constant regressors were included. Condition regressors were convolved with the canonical hemodynamic response function (HRF). A high-pass filter with a cut-off frequency of 128 s was used to remove low-frequency drifts. The collinearity among all regressors of the anticipation, response and outcome phases was modest (*M* = −0.0145; *Median* = −0.0355; *SD* = 0.1074; *Min* = −0.1762; *Max* = 0.6057), consistent with recommended limits (Mumford et al., 2015).

In addition, since the described GLM might conflate anticipation and response phases, we set up a second model to disentangle transient neural activity tied strictly to event onsets (e.g., response initiation or inhibition) from sustained phase-related signals. Therefore, we ran a control GLM with all regressor durations set to 0 (effectively modeling events as instantaneous impulses). This approach isolates activity linked to the precise timing of events rather than their temporal extent. Both models yielded highly consistent findings (see Table 2, Table S9), confirming the robustness of our results.

#### Second level analyses

Second-level GLMs for traditional univariate analyses were estimated based on contrast images from the first-level analyses of all participants and tested against zero in one-sample *t*-tests. To isolate neural activity associated with anticipating affective outcomes, we compared each affective condition to the neutral condition. The contrasts of interest involved appCS+ vs. neuCS−, avCS+ vs. neuCS−, and confCS+ vs. neuCS−. To identify regions distinguishing conflict from pure reward or threat anticipation, we examined the contrasts confCS+ vs. appCS+ and confCS+ vs. avCS+. As the confCS+ was associated with both reward and threat, we further performed conjunction analyses to reveal brain regions commonly activated by confCS+ and/or appCS+/avCS+ (relative to neuCS−) using the minimum statistic approach (Friston et al., 2005). This approach identifies voxels significantly activated in both contrasts of each conjunction, revealing regions engaged across overlapping affective processes. Significant clusters from these conjunctions were used to define regions of interest (ROIs) for subsequent psychophysiological interaction (PPI) and multivariate pattern (MVPA) analyses (see below).

To identify the brain regions involved in overcoming automatic approach-avoidance tendencies, we focused on forced trials and computed contrasts between forced approach (forcedApp) and forced avoidance (forcedAvd) for each CS type. This contrast isolated brain activity associated with executing effortful, counter-motivational actions (e.g., approaching avCS+ vs. avoiding avCS+, and avoiding appCS+ vs. approaching appCS+).

Free trials were excluded from direct approach vs. avoidance contrasts due to low behavioral variance (i.e., insufficient discordant responses; see Behavioral Results). Our primary focus was conflict resolution under externally imposed conditions, as captured in forced trials. Exploratory analyses, including covariates of behavioral metrics (e.g., individual differences in the proportion of avoidance behavior), were conducted but yielded no significant effects (see results at https://osf.io/j5hze).

Imaging analyses are reported at *p* < 0.05, family-wise error (FWE) corrected at the cluster level after an initial cluster-defining threshold of *p* < 0.001 across the whole brain or at *p* < 0.05 small-volume corrected (SVC) at the peak voxel level (FWE-SVC) using anatomical masks from regions of strong a priori interest, derived from the Automated Anatomical Labeling (AAL) atlas (Rolls et al., 2020). These ROIs were defined a priori on anatomical grounds and independently of the present data, including ventral striatum (VS), amygdala, insula, anterior cingulate cortex (ACC), middle cingulate cortex (MCC), inferior frontal gyrus (IFG), and anterior prefrontal cortex (aPFC). These regions were selected based on their theoretical relevance to affective and cognitive processing as described in the introduction. As movement was explicitly required in the AAC task, we also checked the role of the precentral gyrus in overcoming threat-driven avoidance and reward-driven approach. All peaks of activation are reported in MNI coordinates. All fMRI figures were created in MRIcroGL, overlaid on the MNI152 standard brain template.

#### Psychophysiological interaction analyses

To investigate how functional connectivity during anticipation of affective outcomes influences the subsequent behavioral responses and leads to differential approach-avoidance responses between affective stimuli (e.g., avCS+ and confCS+), we performed psychophysiological interaction (PPI) analyses. We used the generalized PPI (gPPI) toolbox (https://www.nitrc.org/projects/gppi), which has the benefit of accommodating multiple task conditions in the same connectivity model (McLaren et al., 2012). Seed regions for connectivity analyses were defined based on the conjunction analyses. Interestingly, only the conjunction of confCS+ > neuCS− ∩ avCS+ > neuCS− revealed significant overlapping activations (Figure 7A), identifying regions engaged during both conflict and threat anticipation. For each seed, a 5-mm spherical ROI was centered on the peak coordinate using MarsBar toolbox (https://www.nitrc.org/projects/marsbar). This led to 5 binary ROIs (Table S11) with 81 voxels in each. Next, we used each ROI as a seed and obtained the respective connectivity strengths with other regions across the brain. In each participant’s first-level PPI model, we included four psychological regressors during anticipation (i.e., 4 task conditions of appCS+, avCS+, confCS+, and neuCS−), one physiological regressor (time series extracted from the seed ROI), and four PPI regressors (i.e., interaction terms, 4 task conditions multiplied by the seed time series, respectively). Besides, 12 nuisance regressors (i.e., head motion parameters and their derivatives, see above) were included.

To identify connectivity differences specific to conflict versus threat anticipation, we contrasted the PPI terms for confCS+ vs. avCS+. Resulting connectivity maps were analyzed at the group level, and significant clusters were thresholded at *p* < 0.05 (cluster-level FWE correction) with a cluster-defining threshold of *p* < 0.001 in the whole-brain analysis. Differences in connectivity strength (confCS+ > avCS+) were correlated with behavioral differences (i.e., ΔP(avoid) in free trials and ΔRT in forced approach and avoidance) between confCS+ and avCS+ using Pearson’s correlation (*p* < 0.05).

#### Multivariate pattern analysis

To examine whether multivariate activation patterns in regions commonly activated by confCS+ and avCS+ (compared to neuCS−) encode information predictive of the subsequent differential behavioral responses, we performed multivariate pattern analysis (MVPA) at the subject level rather than the trial level due to limited trial numbers. This approach is sensitive to distributed spatial patterns that may not be detectable via univariate contrasts. We focused on the confCS+ vs. avCS+ contrast to isolate neural representations specific to conflict anticipation versus pure threat anticipation, despite their overlapping univariate activity. We extracted parameter estimates (beta weights) for each participant for the confCS+ > avCS+ contrast within the conjunction-derived ROIs. These estimates were transformed into N-dimensional pattern vectors (81 voxels per ROI, Table S11). A leave-one-subject-out cross-validated support vector regression (SVR) model was trained to predict subsequent behavioral differences between confCS+ and avCS+ from neural patterns. The SVR was performed by using cosanlab/nltools package (version 0.5.1; https://zenodo.org/records/10888639) with a linear kernel and a preselected cost parameter of 0.01 (Chang et al., 2024). Model performance was evaluated using permutation testing (5000 iterations) with a significance threshold of *p* < 0.05.

To confirm the robustness of our findings based on conjunction-derived ROIs, we additionally defined ROIs using meta-analytic maps from Neurosynth (https://neurosynth.org) based on the terms “dacc” (dorsal ACC), “middle cingulate”, “ventral striatum”, “ifg” (inferior frontal gyrus), and “anterior insula”. For each term, a 5-mm spherical ROI was centered on the peak coordinate from the Neurosynth-derived activation map (Table S11). Analyses using these meta-analytic ROIs yielded largely consistent results (Figure 7E-F, Table S11).

## Results

### Ratings

To examine the affective evaluation of the different CSs, we conducted one-way ANOVAs on valence and arousal ratings (Figure 2A and 2B). These analyses revealed a significant effect of CS type on valence, *F*(3, 117) = 45.11, *p* < 0.001, η_p_^2^ = 0.54, and arousal, *F*(3, 117) = 34.32, *p* < 0.001, η_p_^2^ = 0.47. As expected, the avCS+ was rated significantly more negatively (*M* = 2.60, *SD* = 2.78) and the appCS+ more positively (*M* = 8.62, *SD* = 2.11) than the other CSs. Importantly, the confCS+ (*M* = 5.36, *SD* = 2.04) was rated significantly more positive than the avCS+ (*t*(39) = 5.43, *p* < 0.001, *d* = 1.18, 95% CI [0.47 1.88]) and significantly more negative than the appCS+ (*t*(39) = −7.13, *p* < 0.001, *d* = −1.39, 95% CI [−2.13, −0.65]). Even though the valence of confCS+ was descriptively lower than that of the neuCS−, no significant difference emerged. Interestingly, arousal ratings were comparable between the three affective stimuli (appCS+, avCS+, and confCS+), which were significantly higher than the neuCS− (*t*(39) = 7.33, *p*s < 0.001, *d* = 1.21, 95% CI [0.59, 1.83]). Full post-hoc comparisons are reported in Supplementary Table S2. Similar patterns to CSs were observed in the valence and arousal ratings of USs, except that the confUS was rated as positively as the appUS (see Supplementary Figure S1). Overall, these results suggest that the experimental manipulations worked, and participants successfully acquired the relationships between CSs and USs and evaluated the CSs in line with the learned outcomes.

**Figure 2.**
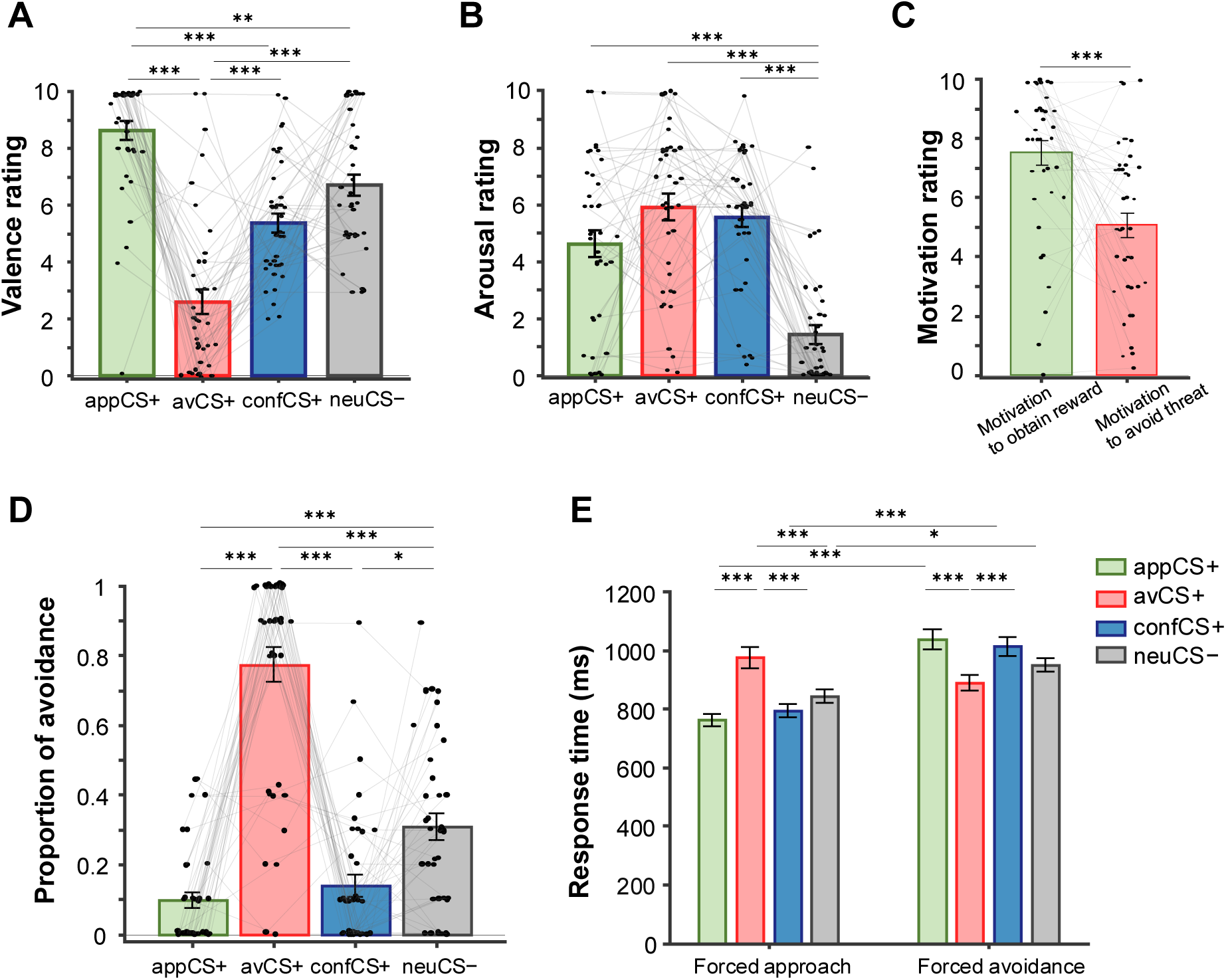
Subjective ratings and behavioral responses in the AAC task. (A-B) Valence and arousal ratings of CSs. (C) Motivation rating to obtain the reward, and to avoid the threat. (D) Proportion of avoidance in free trials. (E) Response times in forced responses. Error bars indicate the standard errors of the mean. * *p* < .05, ** *p* < .01, *** *p* < .001.

Regarding the general motivation to approach reward and avoid threat (Figure 2C), participants on average reported higher motivation to obtain the reward as compared to avoid the aversive electrical stimulation (paired samples *t-*test, *t*(39) = 4.05, *p* < 0.001, *d* = 0.64, 95% CI [0.30 0.98]), suggesting that at the group level, the potential for reward approach was more appealing than the potential for threat avoidance. However, the discrepancy in higher motivation to obtain reward than to avoid threat was not associated with any of the measured personal traits or response choices to the appCS+, avCS+, or confCS+. Intriguingly, individual differences were evident (see the spread of individual data points in Figure 2C), and while many participants followed the general trend of higher motivation for rewards, some exhibited the opposite pattern, showing greater motivation to avoid the aversive stimulation than to obtain the reward. This suggests motivational priorities are not uniform across individuals, especially in conflict situations. Overall, most participants found the task relatively easy (low decision difficulty, *M* = 2.08, *SD* = 1.92).

### Behavioral responses

#### Individual movement trajectories

The individual joystick movement trajectories towards the approach or avoidance options in the free and forced trials for all CSs in the ACC task are illustrated in Figure 3. In the forced trials, participants showed evident delays in initiating joystick movement when required to avoid the reward and approach the threat (Figure 3E and 3F), as quantified by response latencies (Figure S2C). Specifically, participants exhibited significantly longer response latencies in forced avoidance for appCS+ trials as compared to forced approach (*t*(39) = 7.24, *p* < 0.001, *d* = −1.18, 95% CI [0.50, 1.86]). Conversely, participants displayed significantly longer response latencies in forced approach for avCS+ trials compared to forced avoidance (*t*(39) = 4.51, *p* < 0.001, *d* = −0.74, 95% CI [0.15, 1.33]). The same response pattern was found for the confCS+ as for the appCS+. Importantly, no significant difference in response latencies was observed for the neuCS− between forced approach and avoidance. Further statistical details on movement characteristics (e.g., execution time and trajectory length) are reported in the Supplementary Information (Table S3).

**Figure 3.**
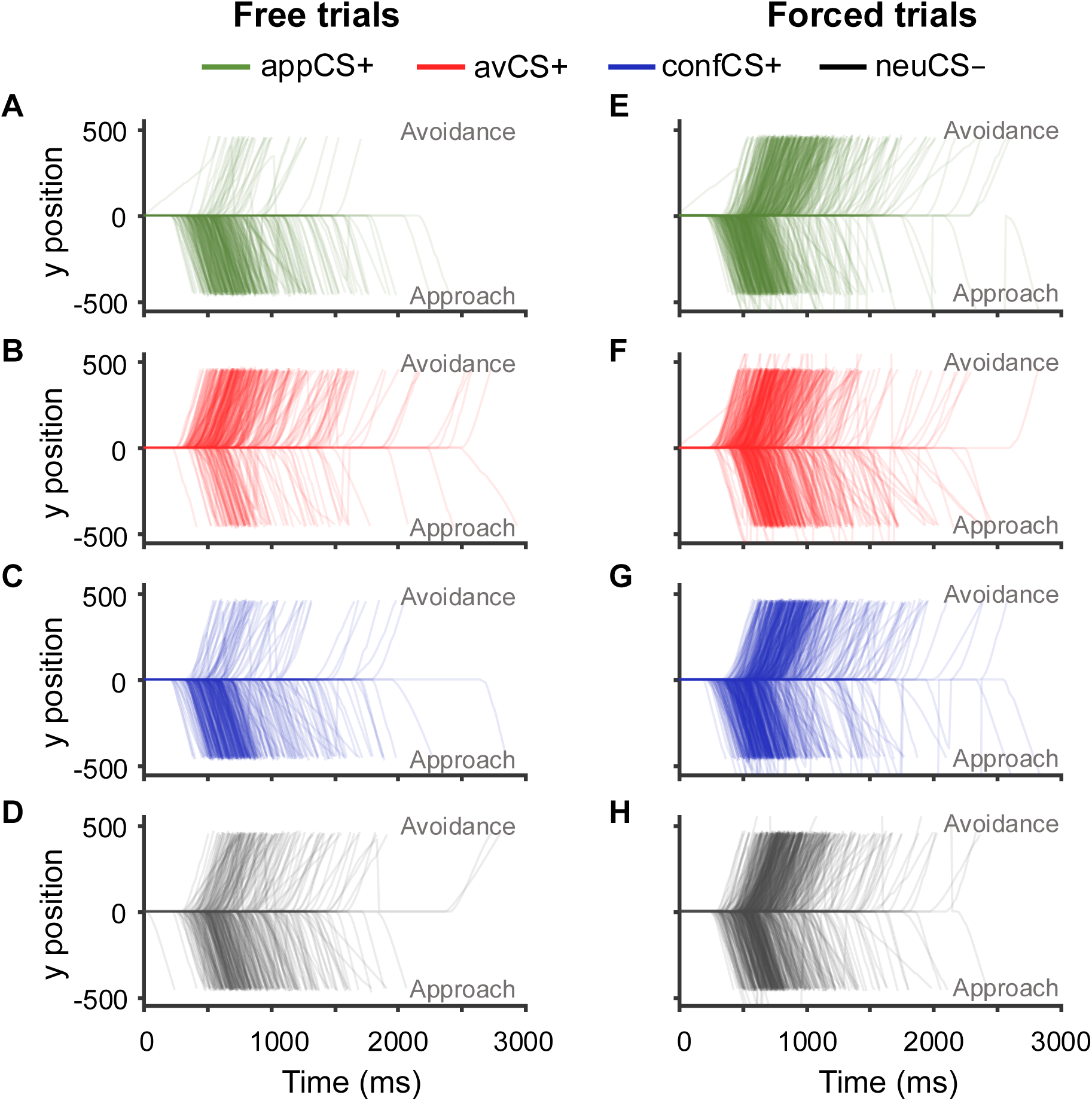
Individual joystick movement trajectories during the free vs. forced trials in the AAC task. In free trials, participants mostly approached the appCS+ (A) and confCS+ (C), and mostly avoided the avCS+ (B). In forced trials, participants showed evident delays in initiating joystick movement when forced to avoid the reward (E) and approach the threat (F). Notably, participants exhibited the same behavioral response pattern for confCS+ and appCS+.

#### Proportion of avoidance in free trials

Results from the one-way ANOVA on the proportion of avoidance (P(avoid), Figure 2D) showed a significant effect of CS type, *F*(3, 117) = 67.18, GG-ε = 0.630, *p* < 0.001, η_p_^2^ = 0.63. As expected, participants showed the least avoidance in the appCS+ condition (*M* = 0.10, *SD* = 0.14) and the most frequent avoidance in the avCS+ condition (*M* = 0.77, *SD* = 0.31). Interestingly, participants demonstrated a strong approach preference for the confCS+ (Figure 2D, 3C). In free trials, the P(avoid) for confCS+ was significantly below chance (*M* = 0.14, *SD* = 0.20; one-sample *t*-test against 0.5: *Z* = −2.29, *p* = 0.022, *d* = −0.36), which was significantly lower than that of the avCS+ (*t*(39) = 11.88, *p* < 0.001, *d* = 2.73, 95% CI [1.70, 3.75]), and the neuCS− (*t*(39) = 3.20, *p* = 0.011, *d* = 0.73, 95% CI [0.08 1.39]). There was no significant difference between the confCS+ and appCS+, suggesting that participants tended to approach despite the aversive electrical stimulation in conflict situations. This behavior matches the affective ratings. Although the confUS was rated as descriptively less positive than the appUS, it was still overall rated highly positive (Figure S1).

#### Response times in forced trials

The 4 × 2 rmANOVA on response times (RTs) in forced trials revealed a significant main effect of response type, *F*(1, 39) = 74.54, *p* < 0.001, η_p_^2^ = 0.18, and a significant interaction of CS type × response type, *F*(3, 117) = 22.33, GG-ε = 0.509, *p* < 0.001, η_p_^2^ = 0.20 (Figure 2E), but no significant main effect of CS type, *F*(3, 117) = 2.05, GG-ε = 0.728, *p* = 0.131, η_p_^2^ = 0.009. As expected, participants showed significantly faster responses to appCS+ in forced approach (*M* = 762.75 ms, *SD* = 133.71 ms) than forced avoidance (*M* = 1036.70 ms, *SD* = 220.29 ms; *t*(39) = −8.38, *p* < 0.001, *d* = −1.54, 95% CI [−2.35 −0.73]). Participants also showed faster forced approach as compared to forced avoidance to neuCS− (approach: *M* = 842.42 ms, *SD* = 142.67 ms; avoidance: *M* = 948.82 ms, *SD* = 139.49 ms; *t*(39) = −3.25, *p* = 0.039, *d* = −0.60, 95% CI [− 1.22 0.03]). Vice versa, participants demonstrated faster responses to avCS+ in forced avoidance (*M* = 890.14 ms, *SD* = 170.34 ms) than forced approach (*M* = 974.82 ms, *SD* = 235.71 ms), which however was not significant (*t*(39) = 2.59, *p* = 0.295, *d* = 0.476, 95% CI [− 0.14 1.09]). Consistent with the P(avoid) variable above, response times for confCS+ followed the same pattern as for appCS+ (Figure 2D-E). For the confCS+, forced approach (*M* = 793.20 ms, *SD* = 148.19 ms) was significantly faster than forced avoidance (*M* = 1013.73 ms, *SD* = 202.32 ms; *t*(39) = −6.74, *p* < 0.001, *d* = −1.24, 95% CI [−1.98 0.50]). Full post hoc tests are provided in Supplementary Table S3.

Taken together, these results indicate that the discordant behavioral responses were initiated and performed more slowly than the concordant responses. Participants more frequently made fast-approach responses to appCS+ and fast-avoidance responses to avCS+. Interestingly, participants also showed an approach preference for confCS+ with faster approach than avoidance responses in forced choices. This mirrors the pattern seen for the appCS+, where approach was dominant, even though the confCS+ was rated more negatively than appCS+.

### Pupillary and oculomotor responses

#### Pupil diameter

The rmANOVA on pupil diameter (Figure 4A) revealed significant main effects of CS type, *F*(3, 105) = 5.83, *p* = 0.001, η_p_ ^2^ = 0.14, and time, *F*(5, 175) = 9.65, GG-ε = 0.252, *p* = 0.002, η_p_^2^ = 0.22, but no significant CS type × time interaction, *F*(15, 525) = 1.46, GG-ε = 0.381, *p* = 0.198, η_p_^2^ = 0.04. These results indicate that both the stimulus condition and time contributed independently to changes in pupil diameter during the later anticipation phase. After the initial pupil constriction following CS onset, pupil diameter increased over time for all conditions, suggesting elevated arousal as the response phase approached. Notably, post-hoc analyses revealed a strong, sustained increase in pupil size in response to stimuli predicting aversive outcomes, with larger dilation for confCS+ than neuCS− (*t*(35) = 4.07, *p =* 0.002, *d* = 0.61, 95% CI [0.14 1.08]), as well as for avCS+ than neuCS− (*t*(35) = 2.52, *p =* 0.033, *d* = 0.36, 95% CI [−0.56 0.77]). No significant differences emerged between appCS+ and neuCS− (*t*(35) = −1.75, *p =* 0.133, *d* = −0.26, 95% CI [−0.68 0.16]). Full post hoc tests are provided in Supplementary Table S4.

**Figure 4.**
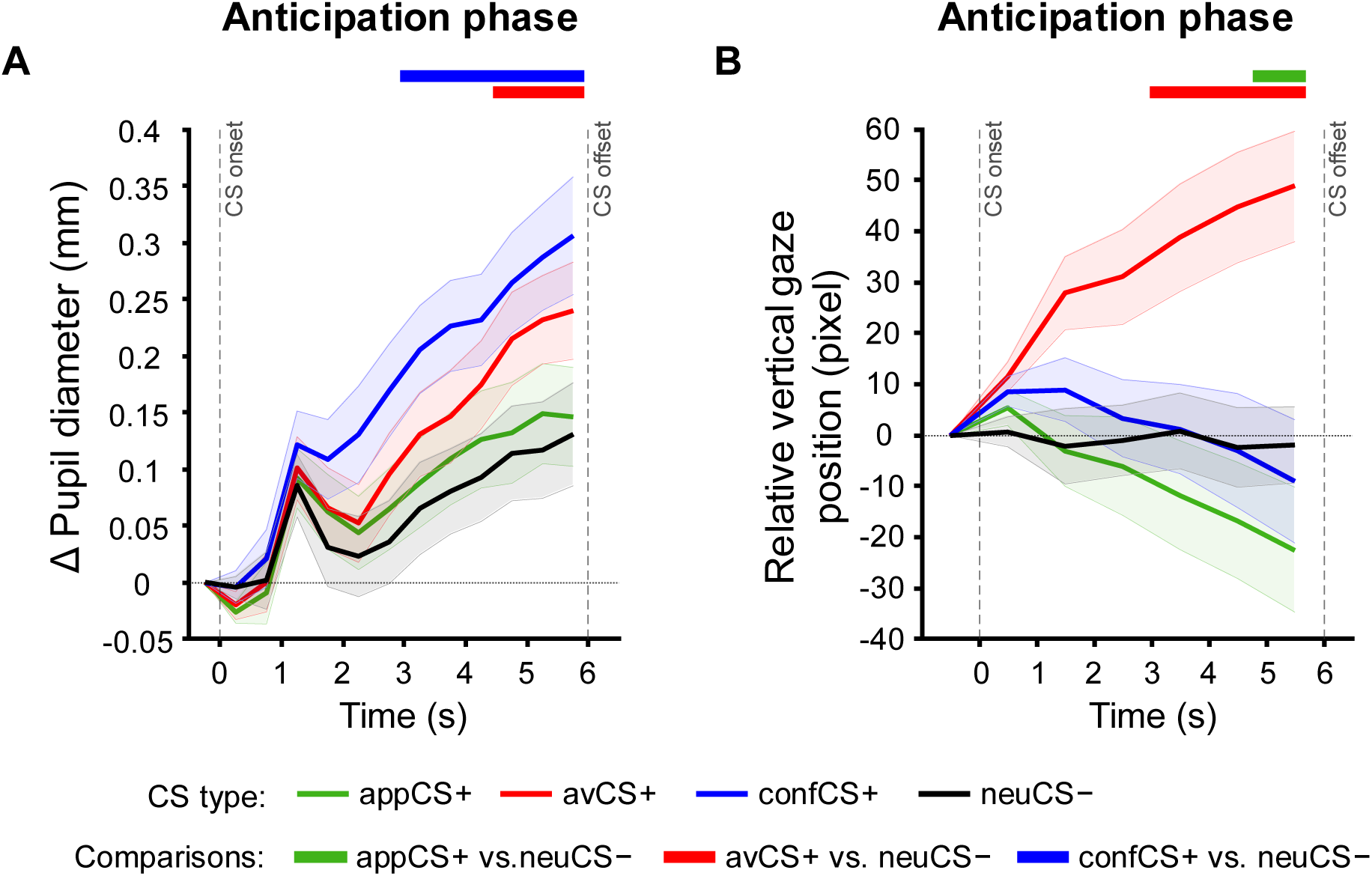
Pupillary and oculomotor responses to CSs during anticipation in the AAC task. (A) Pupil diameter changes. (B) Relative vertical gaze position. Shaded ribbons denote standard errors of the mean. Horizontal lines at the top of the plot indicate significant pairwise differences between the appetitive (appCS+), aversive (avCS+), and conflicting (confCS+) conditions and the neutral CS (neuCS−), respectively.

Moreover, greater pupil dilation during the last second of anticipation was associated with faster approach responses for appCS+ (*r* = −0.37, *p* = 0.029, |Z| = 0.38, 95% CI [−0.62 −0.04]) and avoidance responses for avCS+ (*r* = −0.33, *p* = 0.047, |Z| = 0.35, 95% CI [−0.60 −0.01]), respectively. This relationship was not observed for confCS+, despite it eliciting greater pupil dilation (Table S5). These results suggest that increased physiological arousal shortly before the response phase had a positive effect on action execution of concordant responses.

### Relative vertical gaze position

The rmANOVA on relative vertical gaze position (Figure 4B) revealed a significant main effect of CS type, *F*(3, 105) = 9.76, GG-ε = 0.407, *p* = 0.002, η_p_^2^ = 0.22, and a significant CS type × time interaction, *F*(6, 210) = 3.89, GG-ε = 0.655, *p* = 0.005, η_p_^2^ = 0.10. The main effect of time was not significant, *F*(2, 70) = 0.91, GG-ε = 0.670, *p* = 0.374, η_p_^2^ = 0.025. Pairwise comparisons showed that participants shifted their gaze upwards for the avCS+ (i.e., increased vertical gaze distance from the center of the screen) and downwards for the appCS+ (decreased vertical gaze distance from the center of the screen) relative to the neuCS−. Interestingly, gaze positions for the confCS+ were located between the avCS+ and the appCS+, with no difference from the neuCS−. This pattern is consistent with our previous finding (Chen et al., 2025). However, unlike the previous study conducted in the behavioral lab, where gaze positions for the confCS+ and neuCS− tended to be in the upper half of the screen, we observed that in the fMRI scanner setting, gaze positions for these stimuli remained rather centered over time. Post hoc tests are provided in Supplementary Table S7.

The observed increased vertical gaze distance for the avCS+ was significantly different from that of neuCS− across the final 3 seconds of CS presentation (*t*(35) = 3.23, *p*s ≤ 0.010, *d* = 0.62, 95% CI [−0.14 1.37]). Difference in vertical gaze distance between the appCS+ and neuCS− reached significance in the final second (*t*(35) = −0.34, *p =* 0.039, *d* = −0.87, 95% CI [−0.87 0.20]). No significant differences emerged for confCS+ versus neuCS− (*t*(35) = 0.05, *p*s ≤ 0.960, *d* = 0.006, 95% CI [−0.41 0.42]). These results suggest that fixations during the CS presentation shifted towards the spatial location associated with the concordant response for single-outcome stimuli (i.e., upward for avoidance of avCS+, and downward for approach for appCS+) even though participants did not yet know whether they would be permitted to execute this response. Collectively, these results indicate that stimuli predicting potential aversive electrical stimulation induced higher anticipatory psychophysiological arousal than the neutral stimulus. Moreover, the stimuli guided attention towards the valence-specific concordant options during anticipation, even though their availability was not disclosed at that phase.

### fMRI results

#### Distinct and shared brain activations during anticipation

The univariate GLMs were used to identify brain regions recruited during the anticipation of reward, threat, and motivational conflict relative to a neutral baseline of no outcome (i.e., contrasts appCS+ vs. neuCS−, avCS+ vs. neuCS−, and confCS+ vs. neuCS−). For the appCS+ > neuCS− contrast (reward anticipation), significant activations were observed in regions associated with reward processing, including ACC and bilateral ventral striatum (Figure 5A, Table 1). In addition, significant activation was also observed in MCC (Table 1). In the avCS+ > neuCS− contrast, we found widespread activations across a network of regions previously implicated in threat-processing, including the ACC, bilateral insula, and MCC (Figure 5B, Table 1). In addition, bilateral IFG and ventral striatum were recruited, but not the amygdala. The confCS+ > neuCS− contrast revealed a broadly similar pattern of distributed activations as threat anticipation, including the ACC, bilateral insula, MCC, bilateral IFG, ventral striatum, and additionally the amygdala (Figure 5C, Table 1). Overall, these contrasts revealed brain regions that encode value and salience. Whole-brain cluster-level familywise error-corrected (FWE-corr) results are reported in Table S8. No significant activations were observed for the inverse contrasts (appCS+, avCS+, or confCS+ < neuCS−).

**Figure 5.**
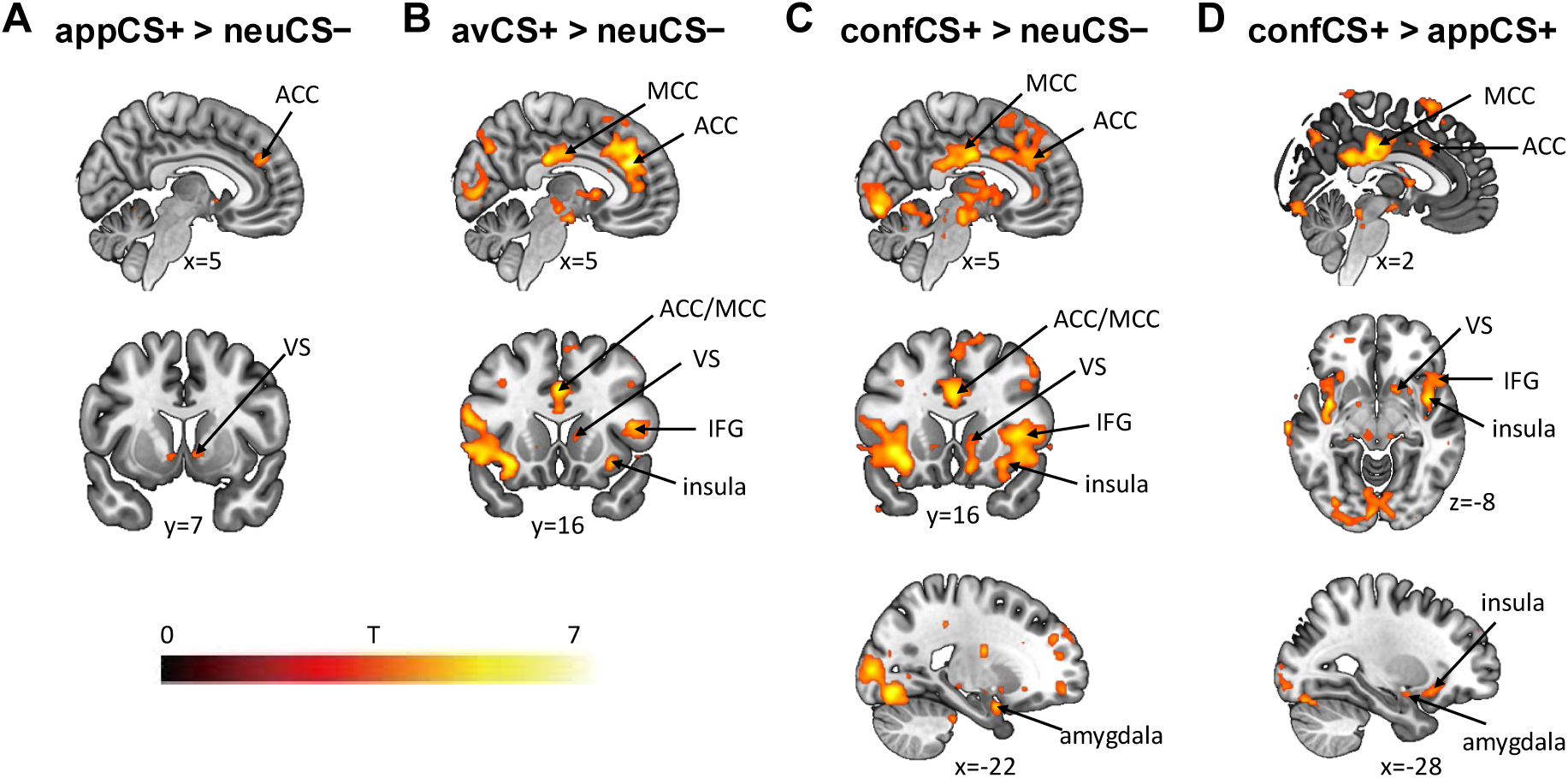
Whole-brain univariate fMRI results demonstrate widespread activations during anticipation of threat and conflict, and a more localized activation during reward anticipation. Images are shown at a threshold of *p* < 0.001 uncorrected. All marked regions of interest reach significance after correction for multiple comparisons (*p*_FWE-SVC_ < 0.05). Abbreviations: anterior cingulate cortex (ACC), middle cingulate cortex (MCC), inferior frontal gyrus (IFG), and ventral striatum (VS).

**Table 1.**
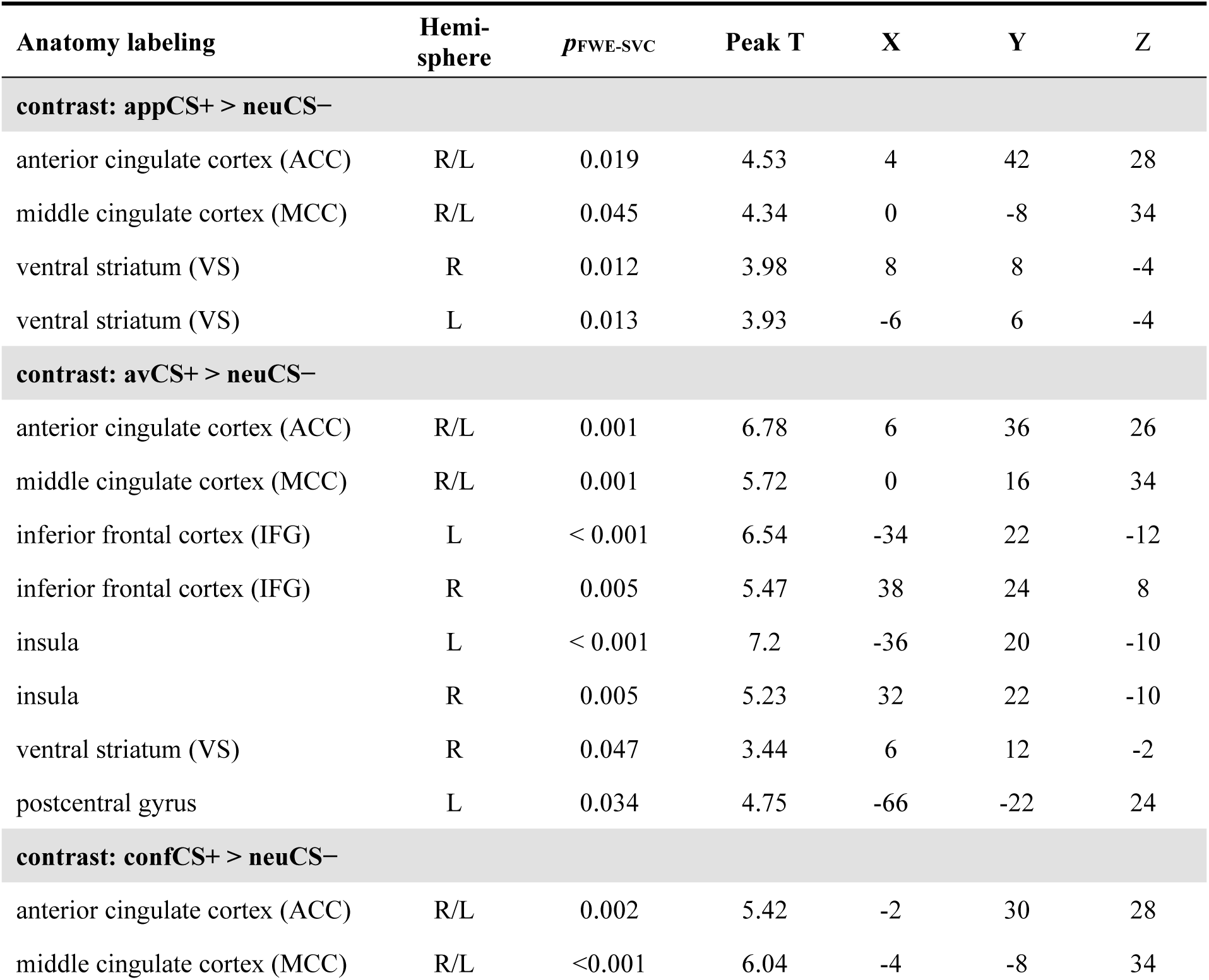
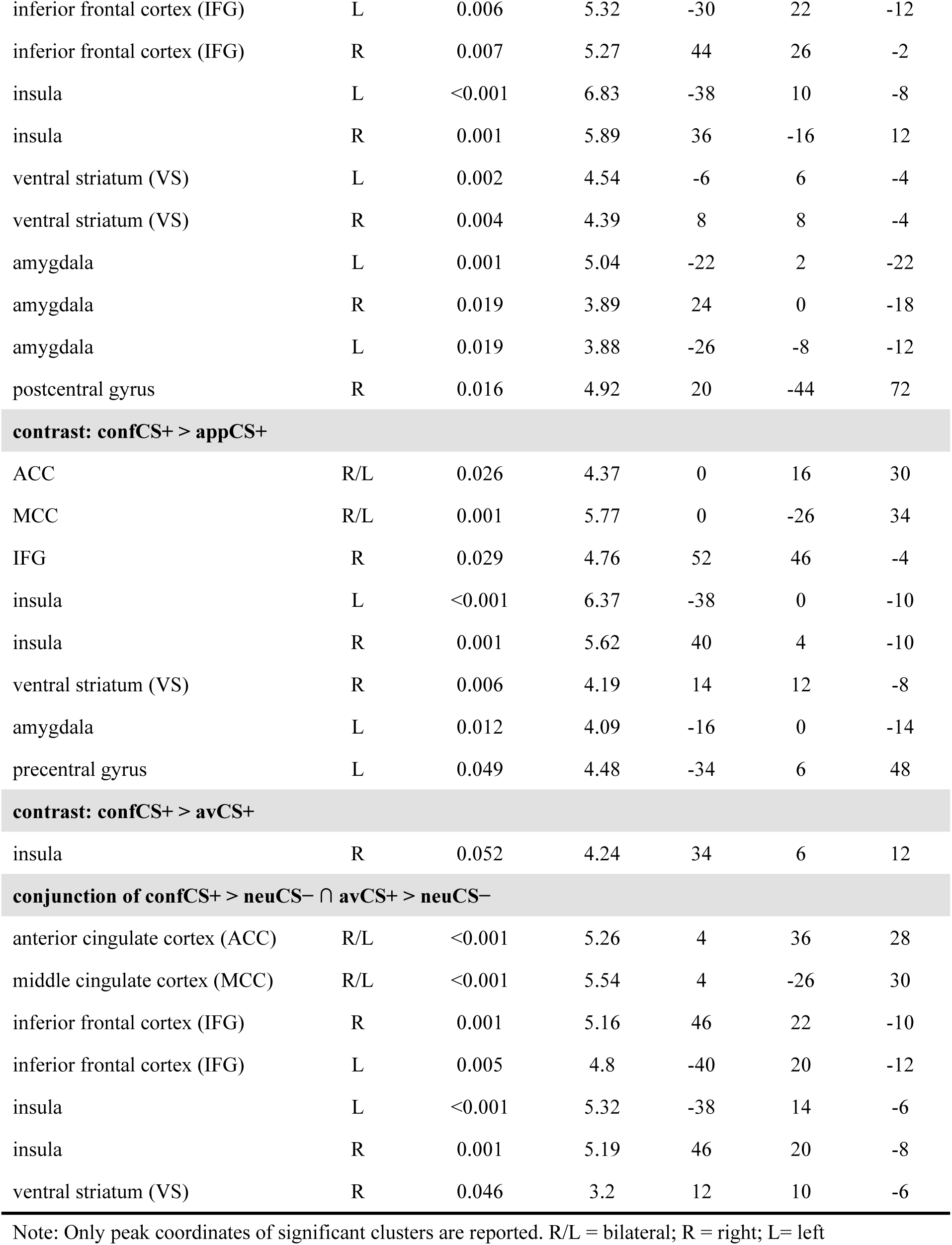
Descriptive statistics of family-wise error-small volume correction (FWE-SVC) for local extrema showing greater activity during the anticipation phase in respective contrast. peak-level *p*_FWE-SVC_ < 0.05.

| Anatomy labeling | Hemi-sphere | $p_{\text{FWE-SVC}}$ | Peak T | X | Y | Z |
| --- | --- | --- | --- | --- | --- | --- |
| **contrast: appCS+ > neuCS-** |  |  |  |  |  |  |
| anterior cingulate cortex (ACC) | R/L | 0.019 | 4.53 | 4 | 42 | 28 |
| middle cingulate cortex (MCC) | R/L | 0.045 | 4.34 | 0 | -8 | 34 |
| ventral striatum (VS) | R | 0.012 | 3.98 | 8 | 8 | -4 |
| ventral striatum (VS) | L | 0.013 | 3.93 | -6 | 6 | -4 |
| **contrast: avCS+ > neuCS-** |  |  |  |  |  |  |
| anterior cingulate cortex (ACC) | R/L | 0.001 | 6.78 | 6 | 36 | 26 |
| middle cingulate cortex (MCC) | R/L | 0.001 | 5.72 | 0 | 16 | 34 |
| inferior frontal cortex (IFG) | L | < 0.001 | 6.54 | -34 | 22 | -12 |
| inferior frontal cortex (IFG) | R | 0.005 | 5.47 | 38 | 24 | 8 |
| insula | L | < 0.001 | 7.2 | -36 | 20 | -10 |
| insula | R | 0.005 | 5.23 | 32 | 22 | -10 |
| ventral striatum (VS) | R | 0.047 | 3.44 | 6 | 12 | -2 |
| postcentral gyrus | L | 0.034 | 4.75 | -66 | -22 | 24 |
| **contrast: confCS+ > neuCS-** |  |  |  |  |  |  |
| anterior cingulate cortex (ACC) | R/L | 0.002 | 5.42 | -2 | 30 | 28 |
| middle cingulate cortex (MCC) | R/L | < 0.001 | 6.04 | -4 | -8 | 34 |
| inferior frontal cortex (IFG) | L | 0.006 | 5.32 | -30 | 22 | -12 |
| inferior frontal cortex (IFG) | R | 0.007 | 5.27 | 44 | 26 | -2 |
| insula | L | <0.001 | 6.83 | -38 | 10 | -8 |
| insula | R | 0.001 | 5.89 | 36 | -16 | 12 |
| ventral striatum (VS) | L | 0.002 | 4.54 | -6 | 6 | -4 |
| ventral striatum (VS) | R | 0.004 | 4.39 | 8 | 8 | -4 |
| amygdala | L | 0.001 | 5.04 | -22 | 2 | -22 |
| amygdala | R | 0.019 | 3.89 | 24 | 0 | -18 |
| amygdala | L | 0.019 | 3.88 | -26 | -8 | -12 |
| postcentral gyrus | R | 0.016 | 4.92 | 20 | -44 | 72 |
| **contrast: confCS+ > appCS+** |  |  |  |  |  |  |
| ACC | R/L | 0.026 | 4.37 | 0 | 16 | 30 |
| MCC | R/L | 0.001 | 5.77 | 0 | -26 | 34 |
| IFG | R | 0.029 | 4.76 | 52 | 46 | -4 |
| insula | L | <0.001 | 6.37 | -38 | 0 | -10 |
| insula | R | 0.001 | 5.62 | 40 | 4 | -10 |
| ventral striatum (VS) | R | 0.006 | 4.19 | 14 | 12 | -8 |
| amygdala | L | 0.012 | 4.09 | -16 | 0 | -14 |
| precentral gyrus | L | 0.049 | 4.48 | -34 | 6 | 48 |
| **contrast: confCS+ > avCS+** |  |  |  |  |  |  |
| insula | R | 0.052 | 4.24 | 34 | 6 | 12 |
| **conjunction of confCS+ > neuCS- ∩ avCS+ > neuCS-** |  |  |  |  |  |  |
| anterior cingulate cortex (ACC) | R/L | <0.001 | 5.26 | 4 | 36 | 28 |
| middle cingulate cortex (MCC) | R/L | <0.001 | 5.54 | 4 | -26 | 30 |
| inferior frontal cortex (IFG) | R | 0.001 | 5.16 | 46 | 22 | -10 |
| inferior frontal cortex (IFG) | L | 0.005 | 4.8 | -40 | 20 | -12 |
| insula | L | <0.001 | 5.32 | -38 | 14 | -6 |
| insula | R | 0.001 | 5.19 | 46 | 20 | -8 |
| ventral striatum (VS) | R | 0.046 | 3.2 | 12 | 10 | -6 |

Note: Only peak coordinates of significant clusters are reported. R/L = bilateral; R = right; L = left

To determine whether regions recruited during threat anticipation exhibit heightened sensitivity to conflict, we directly compared the confCS+ and avCS+. The right insula showed increased activation during conflict anticipation at trend-level significance (*p*_FWE-SVC_ = 0.052, Table 1). However, the direct comparison of confCS+ > appCS+ revealed that conflict processing recruited greater activation in the ACC, MCC, insula, amygdala, IFG, precentral gyrus, and ventral striatum (Figure 5D, Table 1). No significant activations were observed for the inverse contrasts confCS+ < appCS+ or confCS+ < avCS+. Overall, threat anticipation recruited a distributed network of subcortical and cortical regions, indicating that threat information seems more salient than reward information during anticipation. These results suggest that both conflict and threat recruited heightened resources for action preparation.

Conjunction analysis of confCS+ > neuCS− ∩ avCS+ > neuCS− identified overlapping neural activations during both conflict and threat anticipation. Key regions included the ACC, MCC, IFG, insula, and ventral striatum (Figure 7A, Table 1). The amygdala (MNI: [−18, 2, −20], T = 3.23, *p*_FWE-SVC_ = 0.058) showed a trend but did not survive correction. These findings suggest that conflict and threat anticipation recruited overlapping circuits which were associated with different behavioral outcomes (see Figure 2 D-E). To follow up on this surprising finding, these regions were used as seeds for connectivity analyses and ROIs for MVPA (see below). Conjunction of confCS+ > neuCS− ∩ appCS+ > neuCS− did not show any significant activations. Together, these results demonstrate that conflicting and aversive stimuli induced similar univariate activations, with the salience network strongly engaged during threat and conflict anticipation and regulation.

#### Brain activations during the response phase

To identify the brain regions involved in overcoming the initial avoidance tendency to threat and the approach tendency to reward, a contrast for forced approach (forcedApp) vs. forced avoidance (forcedAvd) for each CS type was examined. In the forcedApp > forcedAvd contrast for the avCS+, we found significant activation in left IFG, MCC, insula, ventral striatum, and precentral gyrus (Figure 6A, Table 2), highlighting a distributed neural network engaged during forced approach towards the aversive stimulus. This may explain how current cognitive goals (forced approach to threat) override conflict arising from previous habits (activated avoidance tendency during anticipation). However, no significant aPFC activation was observed. In addition, the contrast forcedAvd > forcedApp for neuCS− revealed significant activation in insula, ventral striatum, and postcentral gyrus (Figure 6B, Table 2). Whole-brain analysis showed significant activations in the postcentral gyrus, insula, cerebellum, superior frontal gyrus, and temporal regions (cluster-level *p*_FWE-corr_ < 0.05, Table S10). These regions have previously been indicated in somatosensory processing, interoceptive awareness, and motor coordination. However, no significant activations were observed when contrasting forcedApp vs. forcedAvd for appCS+ or confCS+.

**Figure 6.**
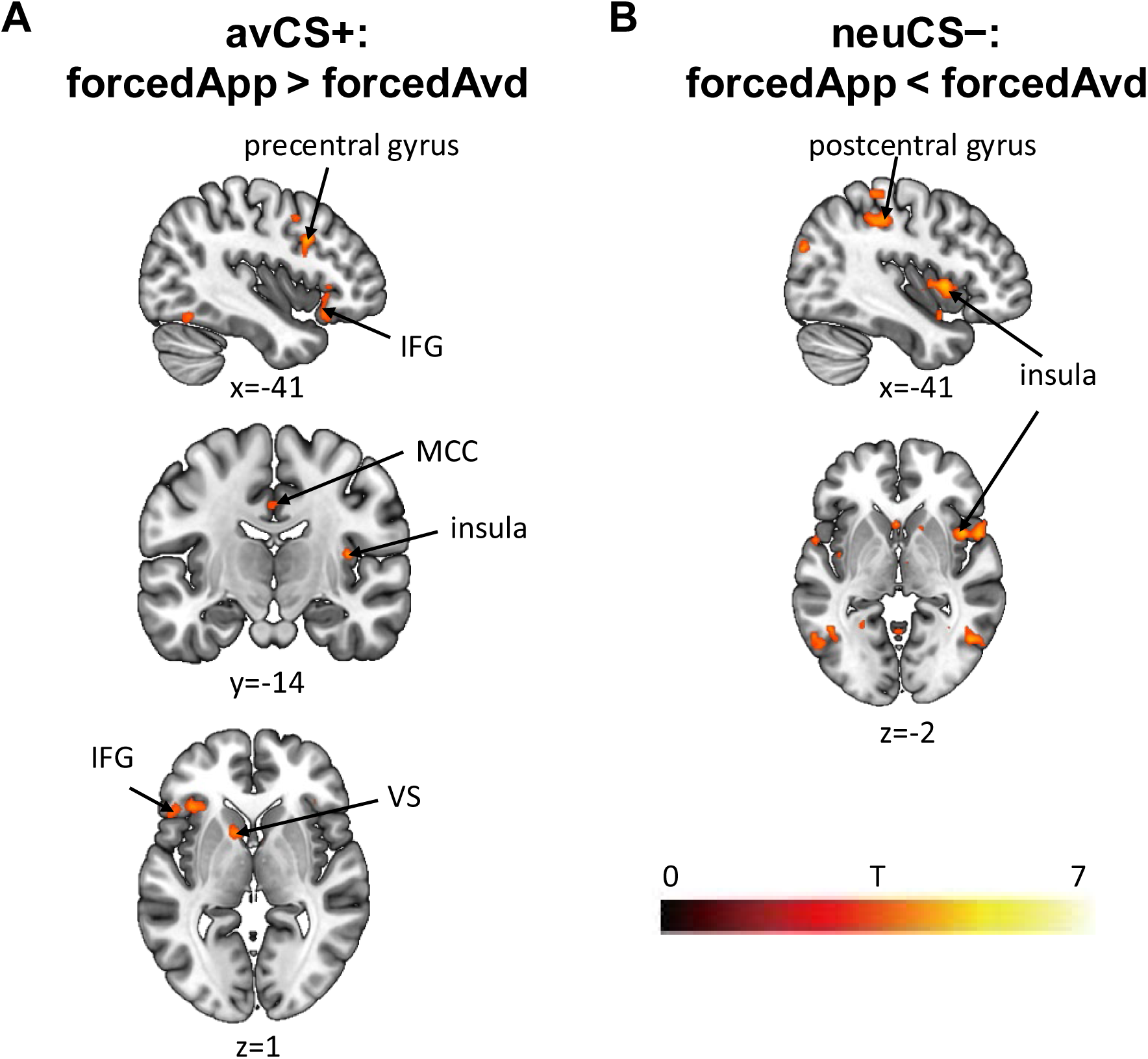
Whole-brain univariate fMRI results of neural activations during response phase. Images are shown at a threshold of *p* < 0.001 uncorrected. All marked regions of interest reach significance after correction for multiple comparisons (*p*_FWE-SVC_ < 0.05). forcedApp = forced approach, forcedAvd = forced avoidance. Abbreviations: middle cingulate cortex (MCC), inferior frontal gyrus (IFG), and ventral striatum (VS).

**Table 2.** Descriptive statistics of family-wise error-small volume correction (FWE-SVC) for local extrema showing greater activity during the response in the respective contrast. peak-level *p*_FWE-SVC_ < 0.05. Durations of response regressors were response times.

| Anatomy labeling | Hemi-sphere | $p_{\text{FWE-SVC}}$ | Peak T | X | Y | Z |
| --- | --- | --- | --- | --- | --- | --- |
| **contrast forcedApp > forcedAvd for avCS+** |  |  |  |  |  |  |
| inferior frontal cortex (IFG) | L | 0.034 | 4.74 | -38 | 22 | -12 |
| middle cingulate cortex (MCC) | L | 0.049 | 4.30 | -12 | -32 | 48 |
| ventral striatum (VS) | L | 0.033 | 3.56 | -10 | 12 | -4 |
| insula | R | 0.016 | 4.74 | 38 | -16 | 12 |
| precentral gyrus | L | 0.020 | 4.87 | -44 | 12 | 32 |
| **contrast forcedApp < forcedAvd for neuCS-** |  |  |  |  |  |  |
| insula | R | 0.019 | 4.73 | 42 | 10 | 2 |
| ventral striatum (VS) | L | 0.024 | 3.73 | -4 | 16 | -2 |
| postcentral gyrus | R | 0.010 | 5.22 | 58 | -18 | 32 |
| postcentral gyrus | L | 0.034 | 4.76 | -48 | -20 | 38 |

Note: Only peak coordinates of significant clusters are reported. R/L = bilateral; R = right; L = left

#### Functional connectivity during anticipation drives behavioral divergence

Despite very similar univariate activation patterns during anticipation of avCS+ (threat) and confCS+ (conflict), participants exhibited divergent behavioral responses: consistent avoidance of avCS+ but approach towards confCS+ (Figure 2D). To investigate whether network interactions distinguish these two conditions, and how functional connectivity during anticipation primes subsequent differential approach-avoidance responses, we conducted PPI analyses using seed regions derived from the conjunction of confCS+ > neuCS− and avCS+ > neuCS− (Figure 7A, Table 1).

**Figure 7.**
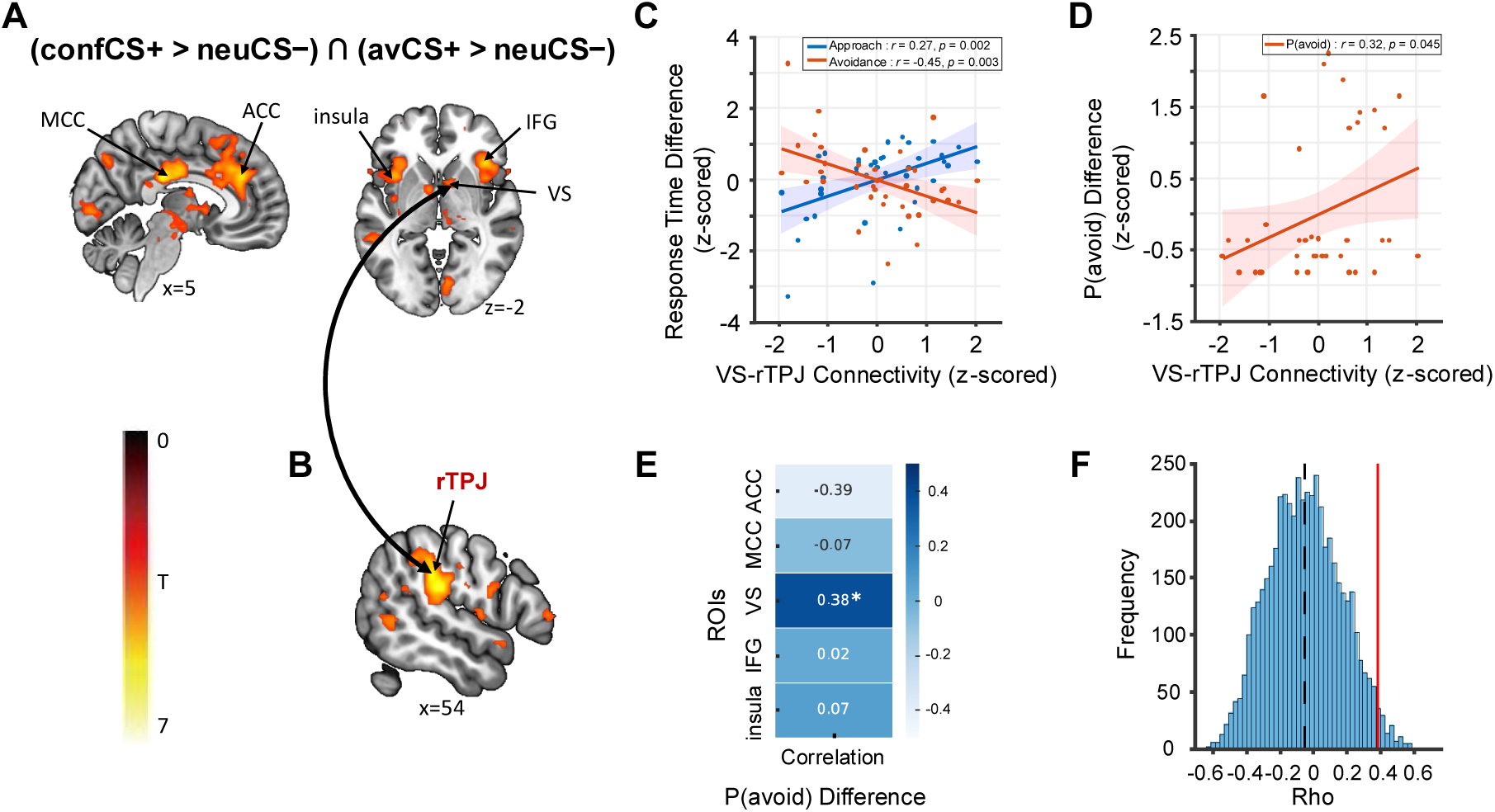
Overlapping univariate neural activations during conflict and threat anticipation, with distinct connectivity and multivariate patterns predicting differential avoidance behavior for avCS+ and confCS+. (A) Conjunction analysis of confCS+ > neuCS− ∩ avCS+ > neuCS− revealed overlapping brain regions commonly activated during both conflict (confCS+) and threat (avCS+) anticipation. These regions served as seeds for psychophysiological interaction (PPI) and multivariate pattern analysis using support vector regression (MVPA-SVR), with a 5-mm spherical region of interest (ROI) centered on the peak coordinate. (B) Whole-brain fMRI results of the PPI analysis demonstrated that enhanced functional connectivity between VS and right temporoparietal junction (rTPJ) was involved in confCS+ compared to avCS+ during anticipation. (C) VS-rTPJ connectivity difference (confCS+ > avCS+) during anticipation significantly correlated with the response time difference between confCS+ and avCS+. (D) VS-rJPG connectivity difference during anticipation significantly correlated with the difference in the P(avoid) in free trials between confCS+ and avCS+. (E) ROI-based neural decoding of P(avoid) differences between confCS+ and avCS+. The heatmap represents the correlation between SVR-predicted and actual values across five ROIs. (F) Permutation testing (K = 5000) confirmed the significance of the VS-based SVR correlation (*r* = 0.38, *p* = 0.023). The black dashed line indicates the mean correlation value from the permutation test, and the red solid line indicates the true correlation value. Images are shown at a threshold of whole-brain analysis *p* < 0.001 uncorrected. All marked regions reach significance after correction for multiple comparisons (*p*_FWE-corr_ < 0.05 or *p*_FWE-SVC_ < 0.05).

Key findings revealed that functional connectivity during anticipation differed between confCS+ and avCS+ and this dissociation correlated with subsequent behavioral divergence. Specifically, during anticipation, connectivity between the VS (seed region) and right temporo-parietal junction (rTPJ, MNI: [54, −26, 22]; k = 1584, *T* = 5.82, cluster-level *p*_FWE-corr_ < 0.001, Figure 7B) was stronger for confCS+ than avCS+. This connectivity was associated with individual differences in subsequent behavioral responses to confCS+ and avCS+ (Figure 7C-D; RTs in forcedAvd, *r* = −0.45, *p* = 0.003; RTs in forcedApp, *r* = 0.47, *p* = 0.002; P(avoid): *r* = 0.32, *p* = 0.045). This suggests individuals with stronger VS-rTPJ connectivity during anticipation for confCS+ (relative to avCS+) exhibited a greater tendency to avoid confCS+ in free trials, as well as faster avoidance but slower approach responses to confCS+ compared to avCS+ in forced trials.

Moreover, enhanced insula-rTPJ coupling (MNI: [48, −36, 34]; k = 413, *T* = 4.50, cluster-level *p*_FWE-corr_ < 0.001) predicted faster avoidance responses to confCS+ relative to avCS+ (RTs in forcedAvd, *r* = −0.39, *p* = 0.012) but did not correlate with approach responses or the proportion of avoidance in free trials. Other seeds (e.g., ACC, IFG) also showed significant connectivity (see Supplementary Table S12) with the rJPJ, but none showed significant behavioral correlations. While ventral striatum and insula also showed significant connectivity with other brain regions like the MCC and the frontal cortex (Table S12), these interactions did not correlate with behavioral differences between confCS+ and avCS+. The MCC seed yielded no significant connectivity. These results identify the right TPJ as a hub integrating signals from the ventral striatum and the insula to shape approach-avoidance decisions, even when univariate activity alone cannot explain the observed behavioral divergence.

#### Activity pattern in the ventral striatum during anticipation encodes behavioral divergence

Participants chose to approach confCS+ (despite its threat component) while avoiding avCS+, even though both showed similar univariate neural activation patterns. Why does confCS+ recruit threat-related regions (like avCS+) but still drive approach behavior (like appCS+)? To address this, we further performed multivariate pattern analysis (MVPA) with support vector regression (SVR) to predict behavioral differences between confCS+ and avCS+ at the subject level. The results showed that neural activity patterns in the ventral striatum during the anticipation phase significantly predicted differences in the proportion of avoidance between confCS+ and avCS+ (*r* = 0.38, permutation testing *p* = 0.023, Figure 7E-F). None of the other conjunction-derived regions (i.e., ACC, MCC, IFG, and insula) showed significant predictive relationships (Figure 7E). These results suggest that multivariate representations in the ventral striatum uniquely encode latent decision variables driving behavioral divergence.

## Discussion

This study examined the neural mechanisms involved in overcoming automatic approach-avoidance tendencies towards affective stimuli (i.e., approaching threat and avoiding reward) and the neural processes engaged in resolving conflicts between opposing action tendencies. The main findings demonstrate that, on a behavioral level, participants exhibited robust concordant responses, i.e., rapid avoidance of threat and approach to reward, accompanied by anticipatory gaze biases toward the respective response-related spatial locations. Responses slowed substantially when required to perform discordant behavior (e.g., approaching threat). These patterns replicate prior work (Chen et al., 2025; Schützwohl, 2018; Volman et al., 2011), and validate the capacity of the task to elicit naturalistic behavioral tendencies while probing inhibitory control.

On a neural level, threat anticipation engaged a broad network involved in salience and threat processing, including the ACC, insula, MCC, IFG, and VS (Ham et al., 2013; Menon & Uddin, 2010; Murty et al., 2023). Crucially, forced approach to threat, relative to forced avoidance, selectively recruited the left IFG (participants used the dominant right hand for joystick movements in the task), consistent with reactive, stimulus-driven inhibition of prepotent avoidance tendencies (Aron et al., 2014; Dignath et al., 2015). In contrast, reward anticipation primarily recruited ACC and VS (Goldstein et al., 2012), and forced reward-avoidance did not elicit differential neural activation relative to reward-approach. A potential interpretation is that suppressing approach to reward may involve devaluing rewards and reducing salience attribution within vmPFC/VS rather than prefrontal inhibition (Bari & Robbins, 2013; Shapcott et al., 2020), which, however, was not captured with the current paradigm. For neutral stimuli, where both approach and avoidance actions led to no reward/punishment, faster forced approach than avoidance recruited sensorimotor regions (postcentral gyrus, cerebellum), consistent with rule compliance without motivational conflict. These findings suggest dissociable substrates for overcoming threat-driven avoidance versus suppressing reward-driven approach. Overall, these patterns support theoretical frameworks of motivated behavior, suggesting that conflicts involving threat elicit stronger cognitive control mechanisms than those involving reward (Jonas et al., 2014; Pessoa, 2009).

Forced approach to threat also engaged a hybrid network associated with action-outcome valuation, such as the MCC (Procyk et al., 2016; Shackman et al., 2011), interoceptive threat signaling implicated in insula regions (Quadt et al., 2018)), and voluntary motor control attributed to the precentral gyrus (Banker & Tadi, 2023). Additionally, ventral striatum activation suggests that approaching threat carried motivational relevance, possibly due to compliance with task demands, e.g., “successfully following instructions” as a form of implicit reward (Deserno et al., 2015). Notably, canonical conflict-resolution regions like the ACC, aPFC, and dorsolateral prefrontal cortex (dlPFC) (Boschin et al., 2017; Botvinick et al., 2001) were not differentially engaged. This may suggest that explicit task instructions, which reduced ambiguity and the need for dlPFC-mediated cognitive control, shifted priorities towards domain-specific inhibition (Egner et al., 2007; Sheu & Courtney, 2016). Our task emphasized rapid, trial-specific inhibition of prepotent responses rather than sustained planning or explicit value comparison. This may explain the predominant reliance on reactive control mechanisms, e.g., IFG (Schaum et al., 2021) rather than dlPFC/aPFC involvement in proactive goal maintenance (Oehrn et al., 2014; Pulopulos et al., 2022).

Interestingly, during anticipation, the conflict cue activated a salience network largely overlapping with threat (ACC, insula, MCC, IFG, VS). Behaviorally, however, participants tended to approach conflict despite high fear responses. Similar behavioral patterns have been reported in previous studies showing that positive outcomes competing with aversive ones elicit approach behavior despite comparable levels of conditioned fear in healthy individuals, whereas this effect seems reduced or absent in individuals with anxiety disorders (Pittig, 2019; Pittig et al., 2020). This divergence, threat-like anticipatory neurophysiology but reward-like approach behavior, suggests that conflict evoked heightened motivational salience and control demands while permitting reward to guide action selection. Psychophysiological interaction analyses revealed stronger VS–right TPJ connectivity during conflict than threat anticipation. Across individuals, higher VS–rTPJ coupling predicted faster avoidance but slower approach to the conflict relative to threat, indicating that this coupling indexes the cognitive cost of reconciling competing valences (Cox & Witten, 2019; Dobryakova & Smith, 2022). The rTPJ, implicated in cognitive flexibility and attentional reorienting, likely facilitates avoidance by dynamically shifting attention toward the threat component in conflict situations (Geng & Vossel, 2013). Within this view, VS-rTPJ coupling may not directly drive approach/avoidance but help regulate the cognitive cost of resolving conflict between them. Stronger connectivity may reflect a ‘careful optimizer’ phenotype (deliberate approach, threat-sensitive decisions), while weaker connectivity may reflect ‘intuitive actors’ (rapid approach, reward-driven choices with reduced threat weighting) (Cohen et al., 2007). Furthermore, VS-rTPJ connectivity might reflect an integration of threat-reward signals with attentional reorienting, enabling a reward-driven override of threat-induced avoidance during conflict. Further studies are warranted to substantiate these hypotheses, validate the suggested phenotypes, and examine their clinical relevance with larger sample sizes.

Further supporting the relevance of the VS in motivated behavior, multivariate analyses revealed decision-relevant information in this structure during the anticipation phase. These patterns predicted individual differences in avoidance between conflict and threat. Analyses using Neurosynth meta-analytic ROIs replicated the conjunction-based results, reinforcing the reliability of our findings. These results indicate that fine-grained striatal representations capture behaviorally relevant motivational states beyond mean univariate activation. This result further proves that motivational and decision-related processes are key to differentiating responses to conflict and threat. Under pure threat, avoidance might be driven by a relatively straightforward Pavlovian fear response (Kim & Jung, 2006; Mobbs et al., 2007), whereas under conflict, avoidance seems to involve a more complex decision-making process, integrating simultaneous reward and threat forecasts, and interfacing with attention/selection systems (e.g., rTPJ) to shape subsequent choice (Aupperle et al., 2015; Dignath et al., 2015; Eder & Hommel, 2013).

Concerning autonomic responses, our results showed that pupil dilation was more pronounced for threat and conflict cues relative to reward and neutral ones. This is consistent with prior work linking pupil dilation to anticipation-related autonomic arousal (Merscher et al., 2022; Skora et al., 2022). Notably, larger pupil dilation at the final second of anticipation was associated with faster concordant responses (approach to reward, avoidance to threat), in line with arousal-facilitated mobilization of action (Aston-Jones & Cohen, 2005). However, this relationship was absent for conflict despite strong dilation, implying that under ambiguity, pupil responses reflect cognitive effort and control demands rather than preparation for a single dominant action (van der Wel & van Steenbergen, 2018; van Steenbergen & Band, 2013). The absence of a behavioral facilitation effect aligns with the competing neural processes observed in our fMRI data (e.g., VS-rTPJ arbitration), where cognitive demand counteracts the typical arousal-action coupling seen in unambiguous conditions.

While the current findings advance our understanding of approach-avoidance conflict processing, several limitations warrant consideration. First, our sample comprised young, healthy, and highly educated participants, limiting generalizability to clinical or demographically diverse populations. Second, although behavioral results replicate prior lab findings (Chen et al., 2025), the fMRI environment may introduce aversive sensations (e.g., scanner noise) that may alter threat processing. Third, participants may have adopted simple heuristic strategies in conflict situations (e.g., default approach in free trials unless forced to avoid), reducing the extent of active deliberation. This is supported by diminished individual variability in conflict responses compared to our prior lab study (Chen et al., 2025), potentially reflecting heightened focus on monetary gains in the effortful fMRI setting. While trial randomization mitigated the predictability of response availability, it may not fully capture naturalistic conflict. Most critically, the competing outcomes differed in modality and timing: pain was immediate and visceral, whereas monetary rewards accumulated over trials and were only paid at the end. This temporal/modality mismatch likely biased conflict salience. Future studies should match outcome modalities and immediacy and incorporate real-time behavioral modeling to disentangle early heuristics from dynamic conflict resolution.

In conclusion, our findings unveiled that aversive stimuli recruited a widespread salience network and elicited strong autonomic responses. Overcoming threat-driven avoidance recruited domain-specific control mechanisms centered on the IFG, whereas suppressing reward-driven approach appeared less effortful and did not elicit increased prefrontal activation. Furthermore, we observed a key dissociation between neural and behavioral signatures of conflict versus threat processing. Although both conditions engaged overlapping neural networks associated with motivational salience, participants consistently avoided threat but approached conflict. Anticipatory VS-rTPJ coupling and multivariate VS patterns predicted individual differences in conflict resolution. Collectively, these findings underscore network-level interactions over isolated regional activity in resolving approach-avoidance conflicts. By dissecting the interplay between automatic and controlled processes, this study advances our understanding of affective control and provide novel insights into neural dynamics of flexible goal-directed behavior.

## Supporting information

https://osf.io/j5hze

## Conflict of interest statement

The authors declare no competing financial interests.

## Funding information

This work was funded by the German Research Foundation (RTG 2660, Project number: 433490190). The funding institution had no role in study design, data collection and analysis, decision to publish, or preparation of the article.

## Acknowledgments

We thank Hannah Zinke and Manuel Danila for their help with data collection, and Prof. Dr. Alexander Shackman and Dr. Jason F. Smith for their insightful advice on fMRI preprocessing and analysis.

## Notes

https://osf.io/j5hze

### Competing Interest Statement

The authors have declared no competing interest.

