## Supplementary material for "Neural Substrates of Approach–Avoidance Control in Motivational Conflict": https://osf.io/j5hze

### Supplementary Information

Supplementary material associated with this article available at the Open Science Framework (https://osf.io/j5hze).

#### Sociodemographic data of participants

**Table S1. Demographic and psychometric data of participants.**

| Variables | Mean (SD) | Range |
|---|---|---|
| Age (years) | 25.15 (4.94) | 19 – 39 |
| Education (years) | 16.93 (3.02) | 11 – 23 |
| Competing reward (Cents) | 9.83 (8.77) | 4 – 30 |
| Electrical stimulation intensity (mA) | 9.19 (8.21) | 0.7 – 46 |
| **General habits** | | |
| Cups of coffee per day | 0.97 (1.04) | 0 – 5 |
| Glasses of alcohol per week | 1.26 (2.55) | 0 – 15 |
| Cigarettes per day | 0.78 (2.39) | 0 – 10 |
| Hours of physical activity per week | 4.11 (4.10) | 0 – 18 |
| **Behavioral Inhibition (BIS) / Approach System Scale (BAS)** | | |
| BIS | 19.75 (4.14) | 17 – 27 |
| BAS Drive | 12.13 (2.22) | 7 – 16 |
| BAS Fun Seeking | 12.35 (2.03) | 8 – 16 |
| BAS Reward Responsiveness | 16.10 (2.35) | 9 – 20 |
| **Depression, Anxiety, and Stress Scale (DASS-21)** | | |
| Depression (DASS-D) | 5.40 (5.01) | 0 – 20 |
| Anxiety (DASS-A) | 4.85 (4.34) | 0 – 16 |
| Stress (DASS-S) | 9.65 (5.56) | 0 – 24 |
| **Big-Five-Inventory-10 (BFI-10)** | | |
| Extraversion | 5.53 (1.93) | 2 – 9 |
| Agreeableness | 5.48 (1.77) | 2 – 9 |
| Conscientiousness | 5.55 (1.74) | 2 – 9 |
| Neuroticism | 4.68 (1.83) | 1 – 8 |

| | | |
|---|---|---|
| Openness | 6.40 (1.81) | 2 – 9 |
| **Decision Making Style** | | |
| Rational | 20.18 (2.70) | 14 – 25 |
| Intuitive | 17.50 (2.76) | 10 – 22 |
| Dependent | 17.20 (3.26) | 6 – 22 |
| Avoidant | 15.28 (4.56) | 6 – 23 |
| Spontaneous | 14.15 (3.67) | 8 – 23 |
| **Intolerance of Uncertainty (IU-18)** | 39.90 (11.63) | 21 – 73 |

Note. SD = standard deviation.

#### Ratings

The one-way ANOVA on valence and arousal ratings of USs (Figure S1A-B) revealed a significant main effect of US type on valence, $F(3, 63) = 69.86$, GG-ε = 0.878, $p < 0.001$, $\eta_p^2 = 0.77$, and arousal, $F(3, 63) = 33.08$, GG-ε = 0.935, $p < 0.001$, $\eta_p^2 = 0.61$. Interestingly, the confUS was rated as positive and arousing as the appUS. Post hoc pairwise comparisons are provided in Table S2. Please note that we only had $n = 22$ participants with ratings data of USs due to technical issues.

For contingency ratings, each CS was presented with the questions, "How likely do you think this shape is followed by the US (i.e., the money, electrical stimulation, both the money and the electrical stimulation, or nothing, respectively)?" The four questions were presented in a pseudo-randomized order for all CSs. The one-way ANOVA revealed that participants demonstrated significantly higher contingency ratings of the paired CS-US associations (i.e., contingency ratings for the correct pair of CS and outcome) than those of unpaired CS-US associations (i.e., contingency ratings for the incorrect outcome) after the AAC task (Figure S1C; appCS+, $F(3, 117) = 234.27$, GG-ε = 0.405, $p < 0.001$, $\eta_p^2 = 0.86$; avCS+, $F(3, 117) = 182.47$, GG-ε = 0.705, $p < 0.001$, $\eta_p^2 = 0.82$; confCS+, $F(3, 117) = 96.81$, $p < 0.001$, $\eta_p^2 = 0.71$; neuCS−, $F(3, 117) = 233$, GG-ε = 0.404, $p < 0.001$, $\eta_p^2 = 0.86$). The contingency awareness confirmed that participants successfully learned the CS-US associations in the AAC task and were able to explicitly report them afterwards.

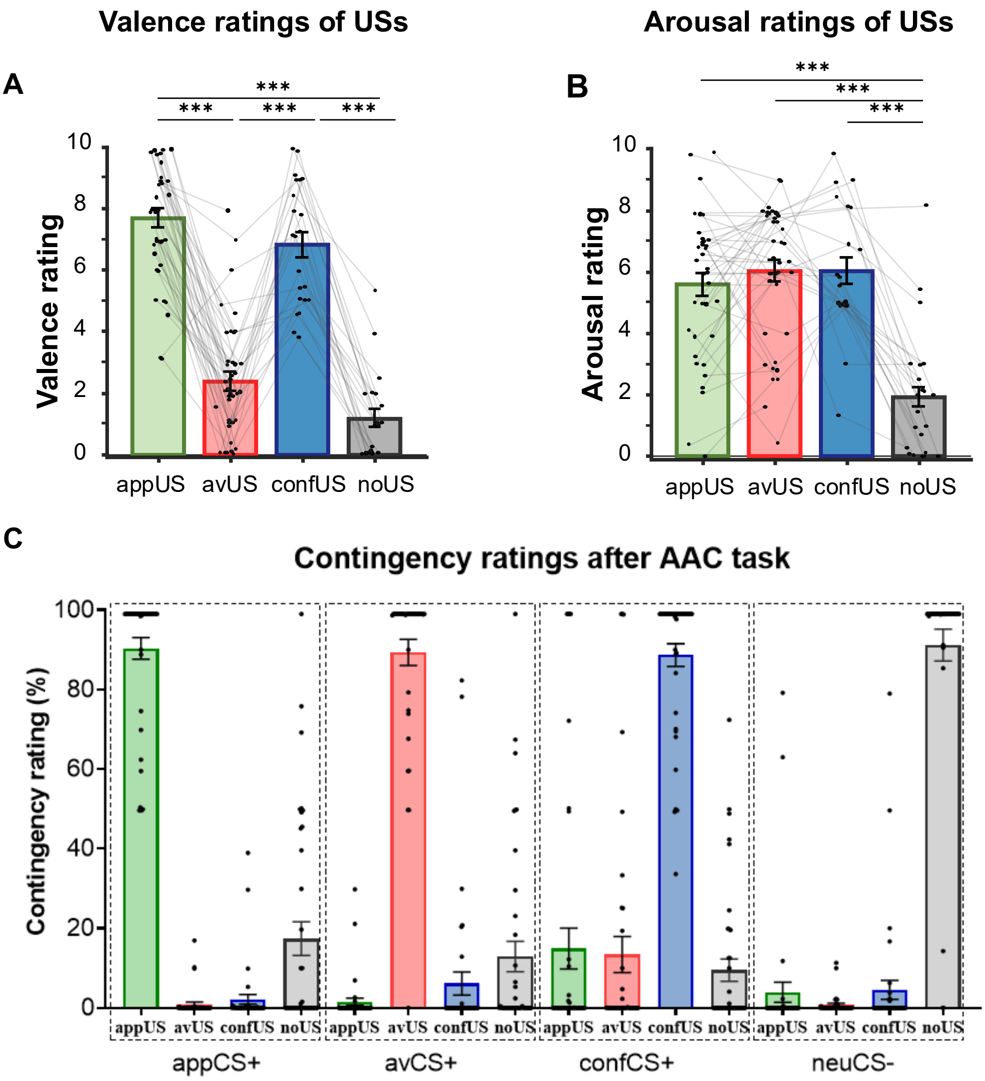


Figure S1. Subjective ratings after the AAC task. (A) Valence ratings of USs. (B) Arousal rating of USs. (C) Contingency ratings. Abbreviations: appUS = appetitive unconditioned stimulus (i.e., monetary reward), avUS = aversive unconditioned stimulus (i.e., electrical stimulation), confUS = conflicting unconditioned stimulus (i.e., both the reward and electrical stimulation), noUS = no outcome.

#### Supplementary Table S2

**Post Hoc Comparisons on ratings across CSs and USs**

| | | | | | | 95% CI for Cohen's *d* | | |
|---|---|---|---|---|---|---|---|---|
| | | **Mean Difference** | **SE** | ***t*** | **Cohen's *d*** | **Lower** | **Upper** | $p_{bonf}$ |
| **Valence of CSs** | | | | | | | | |
| appCS+ | avCS+ | 6.02 | 0.631 | 9.54 | 2.564 | 1.58 | 3.549 | < .001 |
| | confCS+ | 3.26 | 0.457 | 7.13 | 1.388 | 0.651 | 2.126 | < .001 |
| | neuCS− | 1.924 | 0.466 | 4.128 | 0.82 | 0.165 | 1.474 | 0.001 |
| avCS+ | confCS+ | -2.761 | 0.509 | -5.429 | -1.176 | -1.879 | -0.473 | < .001 |
| | neuCS− | -4.096 | 0.584 | -7.016 | -1.745 | -2.548 | -0.941 | < .001 |
| confCS+ | neuCS− | -1.335 | 0.527 | -2.534 | -0.569 | -1.199 | 0.061 | 0.092 |

| | | | | | | | | |
|---|---|---|---|---|---|---|---|---|
| **Arousal of CSs** | | | | | | | | |
| appCS+ | avCS+ | -1.292 | 0.589 | -2.192 | -0.489 | -1.012 | 0.033 | 0.206 |
| | confCS+ | -0.95 | 0.423 | -2.245 | -0.36 | -0.873 | 0.153 | 0.183 |
| | neuCS− | 3.188 | 0.435 | 7.327 | 1.208 | 0.587 | 1.829 | < .001 |
| avCS+ | confCS+ | 0.342 | 0.485 | 0.704 | 0.13 | -0.373 | 0.632 | 1 |
| | neuCS− | 4.479 | 0.556 | 8.06 | 1.698 | 0.979 | 2.416 | < .001 |
| confCS+ | neuCS− | 4.138 | 0.445 | 9.298 | 1.568 | 0.877 | 2.259 | < .001 |
| **Valence of USs** | | | | | | | | |
| appUS | avUS | 5.635 | 0.667 | 8.452 | 3.067 | 1.545 | 4.588 | < .001 |
| | confUS | 1.097 | 0.572 | 1.918 | 0.597 | -0.265 | 1.459 | 0.413 |
| | noUS | 6.734 | 0.529 | 12.717 | 3.665 | 1.928 | 5.401 | < .001 |
| avUS | confUS | -4.538 | 0.617 | -7.35 | -2.47 | -3.789 | -1.15 | < .001 |
| | noUS | 1.099 | 0.506 | 2.172 | 0.598 | -0.264 | 1.46 | 0.249 |
| confUS | noUS | 5.637 | 0.446 | 12.649 | 3.068 | 1.546 | 4.59 | < .001 |
| **Arousal of USs** | | | | | | | | |
| appUS | avUS | 0.163 | 0.443 | 0.367 | 0.078 | -0.619 | 0.775 | 1 |
| | confUS | 0.431 | 0.584 | 0.739 | 0.207 | -0.494 | 0.909 | 1 |
| | noUS | 4.518 | 0.555 | 8.139 | 2.172 | 1.025 | 3.318 | < .001 |
| avUS | confUS | 0.269 | 0.541 | 0.497 | 0.129 | -0.569 | 0.827 | 1 |
| | noUS | 4.355 | 0.527 | 8.266 | 2.093 | 0.973 | 3.214 | < .001 |
| confUS | noUS | 4.086 | 0.537 | 7.607 | 1.964 | 0.886 | 3.043 | < .001 |

Note. For each variable, P-value and confidence intervals were corrected using the Bonferroni method. Abbreviations: appCS+ = appetitive CS+, avCS+ = aversive CS+, confCS+ = conflicting CS+, neuCS− = neutral CS-, appUS = appetitive US, avUS = aversive US, confUS = conflicting US, noUS = no US (i.e., the corresponding CS was neutral).

#### Movement characteristics during the AAC task

The omission rate was calculated as the number of missed trials in each condition divided by the total trials (i.e., 10 free trials or 20 forced trials for each CS, respectively). The rmANOVA revealed that participants missed more trials of avCS+ in forced approach than the other three CSs, $F(3, 117) = 11.82$, GG-$\varepsilon = 0.384$, $p < 0.001$, $\eta_p^2 = 0.23$. A potential explanation is that participants deliberately missed these trials as a strategy to get rid of the aversive electrical stimulation, despite being aware of the associated cost of reduced bonus.

We extracted additional variables based on joystick movement trajectories, including response latency, execution time, and trajectory length. Response latency was defined as the time interval between the manikin onset and first initiation of the joystick movement. Execution time was calculated as the time between the first initiation of joystick movement and the manikin's arrival at the available open-door area. Trajectory length in unit of pixel was the sum of the Euclidean distances between adjacent joystick positions in each trial.

The rmANOVA on these variables revealed a significant main effect of response type (response latency: $F(1, 39) = 38.91$, $p < 0.001$, $\eta_p^2 = 0.50$; execution time: $F(1, 39) = 33.29$, $p < 0.001$, $\eta_p^2 = 0.46$; trajectory length, $F(3, 39) = 9.69$, $p = 0.003$, $\eta_p^2 = 0.20$), and a significant interaction between CS type and response type (response latency: $F(3, 117) = 25.92$, GG-$\varepsilon = 0.532$, $p < 0.001$, $\eta_p^2 = 0.40$; execution time: $F(3, 117) = 3.74$, GG-$\varepsilon = 0.570$, $p = 0.035$, $\eta_p^2 = 0.09$; trajectory length, $F(3, 117) = 4.022$, GG-$\varepsilon = 0.384$, $p = 0.033$, $\eta_p^2 = 0.09$). Post hoc *t*-tests are provided in Table S3.

Pairwise comparisons showed that participants exhibited significantly longer response latencies when required to perform discordant responses (i.e., forced avoidance to the appCS+ and approach to the avCS+). Response latency did not differ between confCS+ and appCS+. Importantly, no significant difference in response latency was observed for the neuCS− between forced approach and avoidance. Regarding execution time, there were no significant differences between forced approach and avoidance for both avCS+ and neuCS−. However, participants exhibited significantly faster execution during forced approach compared to forced avoidance for both appCS+ and confCS+. For trajectory length, a significant difference emerged only for appCS+, with participants showing longer trajectories during forced avoidance to appCS+ compared to forced approach. These findings align with the response times.

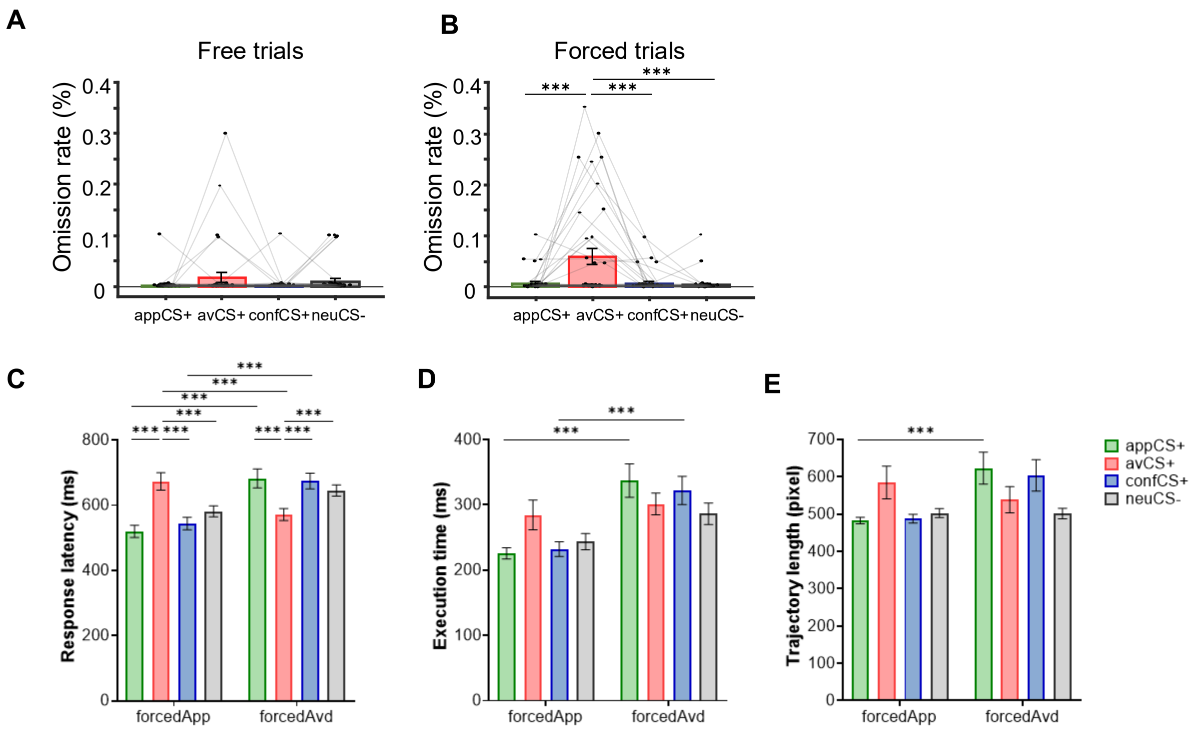


**Figure S2.** Omission rate (A-B) and additional variables based on joystick movements in the AAC task. (C) Response latency, the time interval between the manikin onset and first initiation of the joystick movement; (D) Execution time, the time between the first initiation of joystick movement and the arrival of the manikin at the available door area; (E) Trajectory length, the sum of the Euclidean distances between adjacent joystick positions in each trial. forcedApp = forced approach; forcedAvd = forced avoidance. Error bars indicate the standard errors of the mean. * p < .05, ** p < .01, *** p < .001.

#### Supplementary Table S3

**Post Hoc Comparisons of joystick movement characteristics in the AAC task: CS type × Response type**

| | | Mean Difference | SE | *t*(39) | Cohen's d | 95% CI for Cohen's d Lower | 95% CI for Cohen's d Upper | $p_{bonf}$ |
|---|---|---|---|---|---|---|---|---|
| **Response time** | | | | | | | | |
| appCS+, ForcedApp | avCS+, ForcedApp | -212.07 | 28.937 | -7.329 | -1.191 | -1.865 | -0.516 | < .001 |
| | confCS+, ForcedApp | -30.453 | 28.937 | -1.052 | -0.171 | -0.694 | 0.352 | 1 |
| | neuCS-, ForcedApp | -79.672 | 28.937 | -2.753 | -0.447 | -0.991 | 0.096 | 0.18 |
| | appCS+, ForcedAvd | -273.951 | 32.7 | -8.378 | -1.538 | -2.347 | -0.73 | < .001 |
| avCS+, ForcedApp | confCS+, ForcedApp | 181.617 | 28.937 | 6.276 | 1.02 | 0.383 | 1.657 | < .001 |
| | neuCS-, ForcedApp | 132.398 | 28.937 | 4.575 | 0.744 | 0.159 | 1.328 | < .001 |
| | avCS+, ForcedAvd | 84.678 | 32.7 | 2.59 | 0.476 | -0.136 | 1.087 | 0.295 |
| confCS+, ForcedApp | neuCS-, ForcedApp | -49.219 | 28.937 | -1.701 | -0.276 | -0.805 | 0.252 | 1 |
| | confCS+, ForcedAvd | -220.534 | 32.7 | -6.744 | -1.238 | -1.976 | -0.501 | < .001 |
| neuCS-, ForcedApp | neuCS-, ForcedAvd | -106.407 | 32.7 | -3.254 | -0.598 | -1.223 | 0.027 | 0.039 |

| | | | | | | | | |
|---|---|---|---|---|---|---|---|---|
| appCS+, ForcedAvd | avCS+, ForcedAvd | 146.558 | 28.937 | 5.065 | 0.823 | 0.225 | 1.421 | < .001 |
| | confCS+, ForcedAvd | 22.964 | 28.937 | 0.794 | 0.129 | -0.392 | 0.65 | 1 |
| | neuCS-, ForcedAvd | 87.871 | 28.937 | 3.037 | 0.493 | -0.055 | 1.042 | 0.076 |
| avCS+, ForcedAvd | confCS+, ForcedAvd | -123.594 | 28.937 | -4.271 | -0.694 | -1.271 | -0.118 | < .001 |
| | neuCS-, ForcedAvd | -58.687 | 28.937 | -2.028 | -0.33 | -0.862 | 0.203 | 1 |
| confCS+, ForcedAvd | neuCS-, ForcedAvd | 64.907 | 28.937 | 2.243 | 0.364 | -0.171 | 0.9 | 0.726 |
| **Response latency** | | | | | | | | |
| appCS+, forcedApp | avCS+, forcedApp | -153.062 | 19.205 | -7.97 | -1.111 | -1.717 | -0.506 | < .001 |
| | confCS+, forcedApp | -23.838 | 19.205 | -1.241 | -0.173 | -0.627 | 0.281 | 1 |
| | neuCS-, forcedApp | -61.555 | 19.205 | -3.205 | -0.447 | -0.925 | 0.031 | 0.044 |
| | appCS+, forcedAvd | -162.609 | 22.469 | -7.237 | -1.181 | -1.861 | -0.501 | < .001 |
| avCS+, forcedApp | confCS+, forcedApp | 129.224 | 19.205 | 6.729 | 0.938 | 0.373 | 1.503 | < .001 |
| | neuCS-, forcedApp | 91.507 | 19.205 | 4.765 | 0.664 | 0.154 | 1.175 | < .001 |
| | avCS+, forcedAvd | 101.246 | 22.469 | 4.506 | 0.735 | 0.145 | 1.325 | < .001 |
| confCS+, forcedApp | neuCS-, forcedApp | -37.717 | 19.205 | -1.964 | -0.274 | -0.734 | 0.187 | 1 |
| | confCS+, forcedAvd | -130.531 | 22.469 | -5.809 | -0.948 | -1.577 | -0.319 | < .001 |
| neuCS-, forcedApp | neuCS-, forcedAvd | -63.835 | 22.469 | -2.841 | -0.464 | -1.016 | 0.089 | 0.143 |
| appCS+, forcedAvd | avCS+, forcedAvd | 110.793 | 19.205 | 5.769 | 0.805 | 0.268 | 1.341 | < .001 |
| | confCS+, forcedAvd | 8.24 | 19.205 | 0.429 | 0.06 | -0.39 | 0.51 | 1 |
| | neuCS-, forcedAvd | 37.219 | 19.205 | 1.938 | 0.27 | -0.19 | 0.73 | 1 |
| avCS+, forcedAvd | confCS+, forcedAvd | -102.552 | 19.205 | -5.34 | -0.745 | -1.27 | -0.22 | < .001 |
| | neuCS-, forcedAvd | -73.573 | 19.205 | -3.831 | -0.534 | -1.024 | -0.044 | 0.005 |
| confCS+, forcedAvd | neuCS-, forcedAvd | 28.979 | 19.205 | 1.509 | 0.21 | -0.245 | 0.666 | 1 |
| **Execution time** | | | | | | | | |
| appCS+, forcedApp | avCS+, forcedApp | -58.96 | 19.66 | -2.999 | -0.523 | -1.109 | 0.062 | 0.085 |
| | confCS+, forcedApp | -6.705 | 19.66 | -0.341 | -0.06 | -0.614 | 0.495 | 1 |
| | neuCS-, forcedApp | -18.109 | 19.66 | -0.921 | -0.161 | -0.718 | 0.397 | 1 |
| | appCS+, forcedAvd | -111.558 | 22.423 | -4.975 | -0.99 | -1.716 | -0.264 | < .001 |
| avCS+, forcedApp | confCS+, forcedApp | 52.255 | 19.66 | 2.658 | 0.464 | -0.115 | 1.043 | 0.237 |
| | neuCS-, forcedApp | 40.851 | 19.66 | 2.078 | 0.363 | -0.207 | 0.932 | 1 |
| | avCS+, forcedAvd | -16.762 | 22.423 | -0.748 | -0.149 | -0.783 | 0.486 | 1 |

| | | | | | | | | |
|---|---|---|---|---|---|---|---|---|
| confCS+, forcedApp | neuCS-, forcedApp | -11.404 | 19.66 | -0.58 | -0.101 | -0.657 | 0.454 | 1 |
| | confCS+, forcedAvd | -89.795 | 22.423 | -4.005 | -0.797 | -1.491 | -0.103 | 0.003 |
| neuCS-, forcedApp | neuCS-, forcedAvd | -42.631 | 22.423 | -1.901 | -0.378 | -1.025 | 0.268 | 1 |
| appCS+, forcedAvd | avCS+, forcedAvd | 35.836 | 19.66 | 1.823 | 0.318 | -0.248 | 0.884 | 1 |
| | confCS+, forcedAvd | 15.058 | 19.66 | 0.766 | 0.134 | -0.423 | 0.69 | 1 |
| | neuCS-, forcedAvd | 50.818 | 19.66 | 2.585 | 0.451 | -0.127 | 1.029 | 0.291 |
| avCS+, forcedAvd | confCS+, forcedAvd | -20.778 | 19.66 | -1.057 | -0.184 | -0.743 | 0.374 | 1 |
| | neuCS-, forcedAvd | 14.983 | 19.66 | 0.762 | 0.133 | -0.424 | 0.69 | 1 |
| confCS+, forcedAvd | neuCS-, forcedAvd | 35.76 | 19.66 | 1.819 | 0.317 | -0.249 | 0.884 | 1 |
| **Trajectory length** | | | | | | | | |
| appCS+, forcedApp | avCS+, forcedApp | -102.037 | 39.962 | -2.553 | -0.533 | -1.219 | 0.153 | 0.318 |
| | confCS+, forcedApp | -5.16 | 39.962 | -0.129 | -0.027 | -0.686 | 0.632 | 1 |
| | neuCS-, forcedApp | -20.297 | 39.962 | -0.508 | -0.106 | -0.766 | 0.554 | 1 |
| | appCS+, forcedAvd | -140.438 | 42.425 | -3.31 | -0.734 | -1.481 | 0.014 | 0.033 |
| avCS+, forcedApp | confCS+, forcedApp | 96.877 | 39.962 | 2.424 | 0.506 | -0.177 | 1.19 | 0.452 |
| | neuCS-, forcedApp | 81.74 | 39.962 | 2.045 | 0.427 | -0.25 | 1.104 | 1 |
| | avCS+, forcedAvd | 46.24 | 42.425 | 1.09 | 0.242 | -0.464 | 0.947 | 1 |
| confCS+, forcedApp | neuCS-, forcedApp | -15.137 | 39.962 | -0.379 | -0.079 | -0.739 | 0.581 | 1 |
| | confCS+, forcedAvd | -116.029 | 42.425 | -2.735 | -0.606 | -1.339 | 0.126 | 0.196 |
| neuCS-, forcedApp | neuCS-, forcedAvd | 1.247 | 42.425 | 0.029 | 0.007 | -0.693 | 0.706 | 1 |
| appCS+, forcedAvd | avCS+, forcedAvd | 84.641 | 39.962 | 2.118 | 0.442 | -0.236 | 1.12 | 0.988 |
| | confCS+, forcedAvd | 19.248 | 39.962 | 0.482 | 0.101 | -0.56 | 0.761 | 1 |
| | neuCS-, forcedAvd | 121.388 | 39.962 | 3.038 | 0.634 | -0.063 | 1.331 | 0.075 |
| avCS+, forcedAvd | confCS+, forcedAvd | -65.393 | 39.962 | -1.636 | -0.342 | -1.012 | 0.329 | 1 |
| | neuCS-, forcedAvd | 36.747 | 39.962 | 0.92 | 0.192 | -0.471 | 0.855 | 1 |
| confCS+, forcedAvd | neuCS-, forcedAvd | 102.14 | 39.962 | 2.556 | 0.534 | -0.153 | 1.22 | 0.316 |

Note. For each variable, p-values and confidence intervals were corrected using the Bonferroni method. forcedApp = forced approach. forcedAvd = forced avoidance

### Post Hoc comparisons of pupillary responses

#### Supplementary Table S4

**Post Hoc Comparisons of pupil diameter during the late anticipation phase in the AAC task**

| | | | | | | 95% CI for Cohen's *d* | | |
|---|---|---|---|---|---|---|---|---|
| | | Mean Difference | SE | *t*(35) | Cohen's *d* | Lower | Upper | *p*FDR |
| **within-subject factor of CS type** | | | | | | | | |
| appCS+ | avCS+ | -0.065 | 0.037 | -1.75 | -0.257 | -0.676 | 0.163 | 0.133 |
| | confCS+ | -0.128 | 0.049 | -2.624 | -0.51 | -1.079 | 0.06 | 0.033 |
| | neuCS- | 0.025 | 0.037 | 0.679 | 0.101 | -0.316 | 0.518 | 0.502 |
| avCS+ | confCS+ | -0.063 | 0.044 | -1.452 | -0.253 | -0.747 | 0.241 | 0.186 |
| | neuCS- | 0.09 | 0.036 | 2.516 | 0.358 | -0.057 | 0.773 | 0.033 |
| confCS+ | neuCS- | 0.153 | 0.038 | 4.072 | 0.611 | 0.144 | 1.077 | 0.002 |
| **within-subject factor of Time (the final 3 s of the CS presentation window)** | | | | | | | | |
| T3.5 | T4 | -0.018 | 0.007 | -2.789 | -0.073 | -0.159 | 0.014 | 0.013 |
| | T4.5 | -0.034 | 0.01 | -3.345 | -0.135 | -0.272 | 0.002 | 0.005 |
| | T5 | -0.059 | 0.015 | -4.02 | -0.235 | -0.439 | -0.031 | 0.003 |
| | T5.5 | -0.074 | 0.02 | -3.766 | -0.294 | -0.563 | -0.024 | 0.003 |
| | T6 | -0.083 | 0.027 | -3.14 | -0.332 | -0.687 | 0.024 | 0.007 |
| T4 | T4.5 | -0.016 | 0.006 | -2.656 | -0.062 | -0.14 | 0.015 | 0.016 |
| | T5 | -0.041 | 0.011 | -3.841 | -0.162 | -0.308 | -0.016 | 0.003 |
| | T5.5 | -0.055 | 0.016 | -3.485 | -0.221 | -0.437 | -0.005 | 0.004 |
| | T6 | -0.065 | 0.023 | -2.8 | -0.259 | -0.566 | 0.048 | 0.013 |
| T4.5 | T5 | -0.025 | 0.007 | -3.621 | -0.1 | -0.194 | -0.005 | 0.003 |
| | T5.5 | -0.04 | 0.013 | -3.123 | -0.159 | -0.329 | 0.012 | 0.007 |
| | T6 | -0.049 | 0.02 | -2.518 | -0.197 | -0.453 | 0.06 | 0.021 |
| T5 | T5.5 | -0.015 | 0.007 | -2.051 | -0.059 | -0.152 | 0.034 | 0.055 |
| | T6 | -0.024 | 0.015 | -1.661 | -0.097 | -0.284 | 0.09 | 0.113 |
| T5.5 | T6 | -0.01 | 0.009 | -1.075 | -0.038 | -0.15 | 0.074 | 0.29 |

Note. For CS type, *p*-value and confidence intervals adjusted for comparing a family of 6 estimates using the false discovery rate (FDR) method. Results were averaged over the levels of Time. For Time, *P*-value and confidence intervals adjusted for comparing a family of 15 estimates using FDR method. Results were averaged over the levels of CS type.

**Supplementary Table S5**

**Pearson's correlations between pupil diameter and response times for forced approach and avoidance across CS type**

| PD | RTs | *N* | *r* | *p* | Lower 95% CI | Upper 95% CI | Effect size (Fisher's z) |
|---|---|---|---|---|---|---|---|
| PD at T6 (appCS+) | RTs for forced approach (appCS+) | 36 | -0.365 | 0.029 | -0.619 | -0.041 | -0.382 |
| PD at T6 (appCS+) | RTs for forced avoidance (appCS+) | 36 | -0.035 | 0.838 | -0.36 | 0.297 | -0.035 |
| PD at T6 (avCS+) | RTs for forced approach (avCS+) | 36 | 0.198 | 0.247 | -0.14 | 0.494 | 0.201 |
| PD at T6 (avCS+) | RT for forced avoidance (avCS+) | 36 | -0.334 | 0.047 | -0.597 | -0.006 | -0.347 |
| PD at T6 (confCS+) | RT for forced approach (confCS+) | 36 | -0.074 | 0.668 | -0.393 | 0.261 | -0.074 |
| PD at T6 (confCS+) | RT for forced avoidance (confCS+) | 36 | 0.201 | 0.241 | -0.137 | 0.496 | 0.203 |
| PD at T6 (neuCS−) | RT for forced approach (neuCS−) | 36 | 0.005 | 0.979 | -0.324 | 0.333 | 0.005 |
| PD at T6 (neuCS−) | RT for forced avoidance (neuCS−) | 36 | -0.087 | 0.612 | -0.404 | 0.248 | -0.088 |

*Note*. T6 means the last second of the anticipation. PD = pupil diameter. RTs = response times.

#### Post Hoc comparisons of oculomotor responses

We also calculated the global center bias based on the fixation data, which is defined as the average absolute distance of fixations from the center of the screen in pixels (Merscher et al., 2022). The rmANOVA on global center bias revealed significant main effects of CS type and Time, but no significant CS type × Time interaction (Table S6). The results showed increased visual exploration after CS onset in the AAC task. However, this effect was primarily driven by vertical, rather than horizontal, eye movements (Supplementary Figure S3 and Table S6). This was in line with the task requirement that vertical screen scanning was mainly needed to detect the availability of response options (Chen et al., 2025)**.**

Across all conditions, global center bias rapidly increased within the first 2 seconds after CS onset and continued to rise steadily until CS offset, suggesting heightened visual exploration in response to CS presentation during the AAC task. Post hoc comparisons for CS types did not yield statistically significant pairwise differences. While descriptive trends (e.g., greater center bias for avCS+ vs. neuCS−) were present, the small effect size ($\eta_p^2$ = .09) suggests limited differences between individual CS types. For Time, post hoc comparisons showed significant differences between later time points, with center bias at T6 significantly greater than at T4 ($t(35) = -2.68$, $p = 0.013$, $d = -0.13$, 95% CI [-0.26 -0.002]). These changes in center bias over time were consistent across CS types, further supporting an overall increase in gaze dispersion as stimulus presentation progressed.

Relative horizontal gaze position was calculated using the same procedure as for the vertical coordinate. The rmAONA revealed no significant effects of CS type or Time (Table S6). This indicates that gaze remained horizontally centralized during the anticipation phase, and the observed increase in global center bias was primarily driven by vertical gaze shifts (see main text **Relative vertical gaze position**).

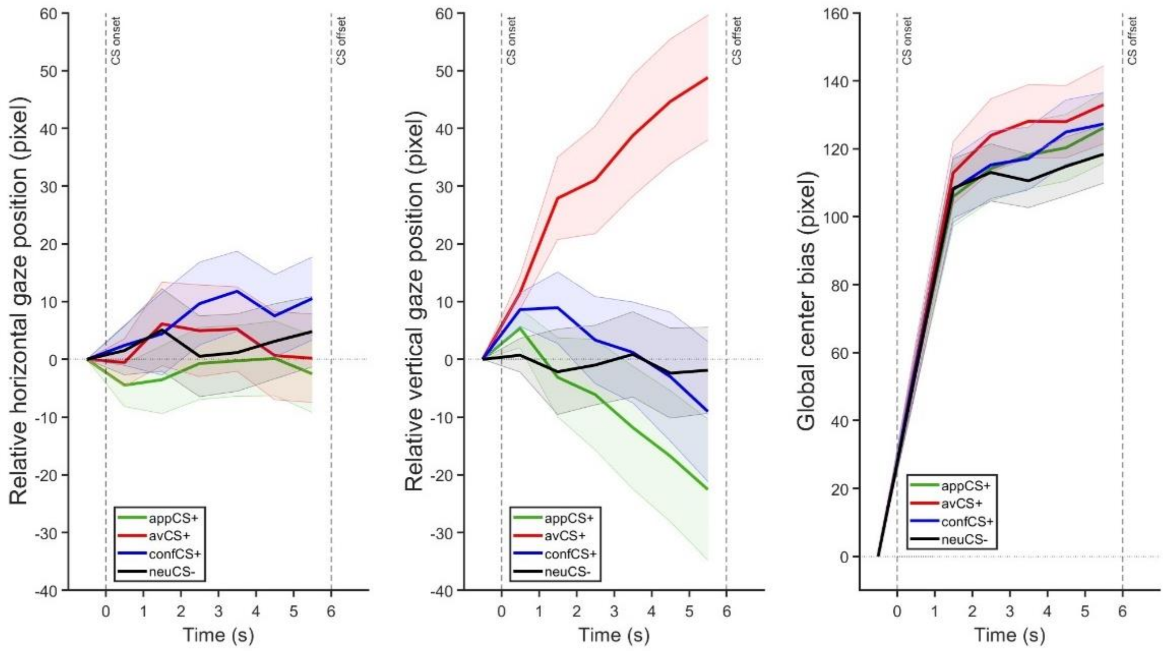


Figure S3. Relative horizontal, vertical gaze position, and global center bias during anticipation in the AAC task.

#### Supplementary Table S6

**Repeated Measures ANOVA on oculomotor responses**

| Items | Greenhouse-Geisser ε | df of factor | df of residuals | Mean Square | *F* | *p* | $\eta_p^2$ |
|---|---|---|---|---|---|---|---|
| **Global center bias** | | | | | | | |
| CS type | 0.779[a] | 3 | 105 | 5318.84 | 3.44 | 0.030 | 0.09 |
| Time | 0.733[a] | 2 | 70 | 2940.04 | 5.74 | 0.011 | 0.14 |
| CS type × Time | 0.686[a] | 6 | 210 | 162.5 | 0.96 | 0.433 | 0.03 |
| **Relative horizontal gaze position** | | | | | | | |
| CS type | 0.711[a] | 3 | 105 | 3192.22 | 1.77 | 0.175 | 0.05 |
| Time | 0.666[a] | 2 | 70 | 154.43 | 0.36 | 0.615 | 0.01 |
| CS type × Time | 0.672[a] | 6 | 210 | 270.26 | 1.73 | 0.147 | 0.05 |

Note. Type III Sum of Squares; [a]Mauchly's test of sphericity indicates that the assumption of sphericity is violated ($p < .05$).

#### Supplementary Table S7

**Post Hoc Comparisons - CS type × Time on vertical gaze position during the late anticipation phase in the AAC task**

| | | Mean Difference | SE | *t*(35) | Cohen's d | 95% CI for Cohen's d Lower | Upper | *p*FDR |
|---|---|---|---|---|---|---|---|---|
| appCS+, T4 | avCS+, T4 | -50.546 | 17.29 | -2.923 | -0.824 | -1.883 | 0.234 | 0.016 |
| | confCS+, T4 | -13.01 | 5.799 | -2.244 | -0.212 | -0.569 | 0.145 | 0.056 |
| | neuCS-, T4 | -12.672 | 7.976 | -1.589 | -0.207 | -0.693 | 0.28 | 0.17 |
| | appCS+, T5 | 4.962 | 2.761 | 1.797 | 0.081 | -0.089 | 0.251 | 0.124 |
| | avCS+, T5 | -56.453 | 17.413 | -3.242 | -0.921 | -1.995 | 0.154 | 0.01 |
| | confCS+, T5 | -8.81 | 6.588 | -1.337 | -0.144 | -0.545 | 0.258 | 0.261 |
| | neuCS-, T5 | -9.417 | 7.709 | -1.222 | -0.154 | -0.622 | 0.315 | 0.297 |
| | appCS+, T6 | 10.744 | 3.907 | 2.75 | 0.175 | -0.072 | 0.423 | 0.022 |
| | avCS+, T6 | -60.607 | 17.902 | -3.386 | -0.989 | -2.098 | 0.121 | 0.009 |
| | confCS+, T6 | -2.773 | 6.913 | -0.401 | -0.045 | -0.462 | 0.371 | 0.735 |
| | neuCS-, T6 | -9.919 | 7.655 | -1.296 | -0.162 | -0.628 | 0.304 | 0.274 |
| avCS+, T4 | confCS+, T4 | 37.535 | 14.408 | 2.605 | 0.612 | -0.296 | 1.52 | 0.029 |
| | neuCS-, T4 | 37.874 | 11.708 | 3.235 | 0.618 | -0.138 | 1.373 | 0.01 |
| | appCS+, T5 | 55.508 | 18.017 | 3.081 | 0.905 | -0.19 | 2.001 | 0.014 |
| | avCS+, T5 | -5.907 | 3.464 | -1.705 | -0.096 | -0.309 | 0.116 | 0.142 |
| | confCS+, T5 | 41.736 | 16.868 | 2.474 | 0.681 | -0.367 | 1.728 | 0.037 |
| | neuCS-, T5 | 41.129 | 12.182 | 3.376 | 0.671 | -0.108 | 1.449 | 0.009 |
| | appCS+, T6 | 61.29 | 18.788 | 3.262 | 1 | -0.137 | 2.136 | 0.01 |
| | avCS+, T6 | -10.061 | 5.182 | -1.941 | -0.164 | -0.484 | 0.156 | 0.1 |
| | confCS+, T6 | 47.773 | 18.24 | 2.619 | 0.779 | -0.339 | 1.898 | 0.029 |
| | neuCS-, T6 | 40.627 | 12.13 | 3.349 | 0.663 | -0.114 | 1.439 | 0.009 |
| confCS+, T4 | neuCS-, T4 | 0.339 | 6.837 | 0.05 | 0.006 | -0.406 | 0.417 | 0.96 |
| | appCS+, T5 | 17.972 | 6.311 | 2.848 | 0.293 | -0.103 | 0.689 | 0.019 |
| | avCS+, T5 | -43.443 | 14.772 | -2.941 | -0.709 | -1.651 | 0.234 | 0.016 |
| | confCS+, T5 | 4.201 | 4.853 | 0.866 | 0.069 | -0.224 | 0.361 | 0.471 |
| | neuCS-, T5 | 3.594 | 6.623 | 0.543 | 0.059 | -0.34 | 0.457 | 0.661 |

| | | | | | | | | |
|---|---|---|---|---|---|---|---|---|
| | appCS+, T6 | 23.755 | 6.806 | 3.49 | 0.387 | -0.05 | 0.825 | 0.008 |
| | avCS+, T6 | -47.597 | 15.604 | -3.05 | -0.776 | -1.776 | 0.223 | 0.014 |
| | confCS+, T6 | 10.237 | 6.314 | 1.621 | 0.167 | -0.217 | 0.551 | 0.164 |
| | neuCS-, T6 | 3.091 | 6.878 | 0.449 | 0.05 | -0.364 | 0.464 | 0.71 |
| neuCS-, T4 | appCS+, T5 | 17.633 | 8.465 | 2.083 | 0.288 | -0.235 | 0.81 | 0.078 |
| | avCS+, T5 | -43.781 | 11.702 | -3.741 | -0.714 | -1.486 | 0.057 | 0.006 |
| | confCS+, T5 | 3.862 | 8.52 | 0.453 | 0.063 | -0.45 | 0.576 | 0.71 |
| | neuCS-, T5 | 3.255 | 2.805 | 1.161 | 0.053 | -0.116 | 0.222 | 0.322 |
| | appCS+, T6 | 23.416 | 9.258 | 2.529 | 0.382 | -0.197 | 0.961 | 0.033 |
| | avCS+, T6 | -47.936 | 11.761 | -4.076 | -0.782 | -1.569 | 0.006 | 0.006 |
| | confCS+, T6 | 9.898 | 9.651 | 1.026 | 0.161 | -0.422 | 0.745 | 0.381 |
| | neuCS-, T6 | 2.753 | 3.746 | 0.735 | 0.045 | -0.181 | 0.271 | 0.532 |
| appCS+, T5 | avCS+, T5 | -61.415 | 18.146 | -3.384 | -1.002 | -2.112 | 0.109 | 0.009 |
| | confCS+, T5 | -13.772 | 5.804 | -2.373 | -0.225 | -0.588 | 0.138 | 0.043 |
| | neuCS-, T5 | -14.378 | 7.966 | -1.805 | -0.235 | -0.725 | 0.256 | 0.124 |
| | appCS+, T6 | 5.782 | 2.897 | 1.996 | 0.094 | -0.085 | 0.274 | 0.091 |
| | avCS+, T6 | -65.569 | 18.616 | -3.522 | -1.069 | -2.213 | 0.074 | 0.008 |
| | confCS+, T6 | -7.735 | 6.17 | -1.254 | -0.126 | -0.502 | 0.249 | 0.288 |
| | neuCS-, T6 | -14.881 | 8.068 | -1.844 | -0.243 | -0.74 | 0.255 | 0.119 |
| avCS+, T5 | confCS+, T5 | 47.643 | 17.096 | 2.787 | 0.777 | -0.295 | 1.849 | 0.021 |
| | neuCS-, T5 | 47.037 | 12.112 | 3.883 | 0.767 | -0.023 | 1.557 | 0.006 |
| | appCS+, T6 | 67.197 | 18.925 | 3.551 | 1.096 | -0.054 | 2.246 | 0.008 |
| | avCS+, T6 | -4.154 | 4.035 | -1.03 | -0.068 | -0.312 | 0.177 | 0.381 |
| | confCS+, T6 | 53.68 | 18.278 | 2.937 | 0.876 | -0.254 | 2.005 | 0.016 |
| | neuCS-, T6 | 46.534 | 12.177 | 3.821 | 0.759 | -0.036 | 1.554 | 0.006 |
| confCS+, T5 | neuCS-, T5 | -0.607 | 7.5 | -0.081 | -0.01 | -0.461 | 0.442 | 0.95 |
| | appCS+, T6 | 19.554 | 5.265 | 3.714 | 0.319 | -0.028 | 0.666 | 0.006 |
| | avCS+, T6 | -51.798 | 17.746 | -2.919 | -0.845 | -1.963 | 0.273 | 0.016 |
| | confCS+, T6 | 6.037 | 3.438 | 1.756 | 0.098 | -0.113 | 0.31 | 0.132 |
| | neuCS-, T6 | -1.109 | 7.636 | -0.145 | -0.018 | -0.478 | 0.442 | 0.913 |
| neuCS-, T5 | appCS+, T6 | 20.161 | 8.322 | 2.423 | 0.329 | -0.193 | 0.85 | 0.039 |

| | | | | | | | | |
|---|---|---|---|---|---|---|---|---|
| | avCS+, T6 | -51.191 | 12.092 | -4.234 | -0.835 | -1.636 | -0.034 | 0.006 |
| | confCS+, T6 | 6.643 | 8.509 | 0.781 | 0.108 | -0.406 | 0.623 | 0.51 |
| | neuCS-, T6 | -0.502 | 2.358 | -0.213 | -0.008 | -0.15 | 0.134 | 0.872 |
| appCS+, T6 | avCS+, T6 | -71.352 | 19.372 | -3.683 | -1.164 | -2.345 | 0.017 | 0.006 |
| | confCS+, T6 | -13.517 | 5.205 | -2.597 | -0.22 | -0.548 | 0.108 | 0.029 |
| | neuCS-, T6 | -20.663 | 8.492 | -2.433 | -0.337 | -0.869 | 0.195 | 0.039 |
| avCS+, T6 | confCS+, T6 | 57.834 | 18.927 | 3.056 | 0.943 | -0.231 | 2.118 | 0.014 |
| | neuCS-, T6 | 50.688 | 12.054 | 4.205 | 0.827 | 0.027 | 1.627 | 0.006 |
| confCS+, T6 | neuCS-, T6 | -7.146 | 8.504 | -0.84 | -0.117 | -0.631 | 0.398 | 0.479 |

Note. P-value and confidence intervals adjusted for comparing a family of 66 estimates using the false discovery rate (FDR) method.

### fMRI results

#### Neural activation during anticipation (Figure 5)

#### Supplementary Table S8

**Descriptive statistics for clusters and local extrema showing greater activity during the anticipation phase in respective contrast at cluster-level $p_{FWE\text{-}corr} < 0.05$ after an initial cluster-defining threshold of $p < 0.001$ across the whole brain.**

| Cluster anatomy (manual labeling) | Hemi-sphere | Cluster size | $p_{FWE\text{-}corr}$ | Peak T | X | Y | Z | Peak label (AAL3) |
|---|---|---|---|---|---|---|---|---|
| **contrast: appCS+ > neuCS−** | | | | | | | | |
| middle cingulate cortex (MCC) | R | 182 | 0.016 | 4.67 | 4 | 42 | 30 | Cingulate_Mid_R |
| **contrast: avCS+ > neuCS−** | | | | | | | | |
| insula | L | 1370 | <0.001 | 7.2 | -36 | 20 | -10 | insula_L |
| anterior cingulate cortex (ACC) | R | 3385 | <0.001 | 6.78 | 6 | 36 | 26 | ACC_sup_R |
| temporo-parietal junction (TPJ) | R | 655 | <0.001 | 6.33 | 62 | -48 | 38 | Supramarginal_R |
| substantia nigra (SN) | R | 228 | 0.003 | 5.78 | 8 | -8 | -16 | SN_pr_R |
| temporo-parietal junction (TPJ) | L | 523 | <0.001 | 5.54 | -62 | -46 | 36 | Supramarginal_L |
| inferior frontal gyrus (IFG) | R | 808 | <0.001 | 5.47 | 38 | 24 | 8 | Frontal_Inf_Tri_R |
| primary visual cortex (V1) | R | 938 | <0.001 | 5.34 | 14 | -78 | 8 | Calcarine_R |
| ventral striatum | R | 245 | 0.002 | 5.1 | 8 | 10 | 4 | Caudate_R |
| visual association cortex | L | 137 | 0.037 | 4.75 | -18 | -94 | 10 | Occipital_Mid_L |
| precuneus | L | 430 | <0.001 | 4.66 | -6 | -70 | 32 | Precuneus_L |
| medial prefrontal cortex (mPFC) | R | 160 | 0.019 | 4.42 | 10 | 58 | 38 | Frontal_Sup_Medial_R |
| middle temporal gyrus (MTG) | R | 143 | 0.031 | 4.32 | 50 | -30 | -6 | Temporal_Mid_R |
| **contrast: confCS+ > neuCS−** | | | | | | | | |
| visual association cortex | R | 4769 | <0.001 | 7.05 | 18 | -92 | 6 | Occipital_Sup_R |
| insula | L | 2943 | <0.001 | 6.83 | -38 | 10 | -8 | Insula_L |
| middle cingulate cortex (MCC) | L | 3879 | <0.001 | 6.04 | -4 | -8 | 34 | Cingulate_Mid_L |
| putamen | R | 2865 | <0.001 | 5.99 | 32 | 6 | 10 | Putamen_R |
| thalamus | L | 1924 | <0.001 | 5.9 | -10 | 0 | 4 | Thal_VA_L |
| insula | R | 219 | 0.015 | 5.89 | 36 | -16 | 12 | Insula_R |
| inferior parietal lobule (IPL) | L | 804 | <0.001 | 4.99 | -40 | -52 | 38 | Parietal_Inf_L |
| postcentral gyrus | R | 165 | 0.048 | 4.92 | 20 | -44 | 72 | Postcentral_R |
| temporo-parietal junction (TPJ) | R | 612 | <0.001 | 4.7 | 52 | -46 | 42 | SupraMarginal_R |
| middle temporal gyrus (MTG) | R | 488 | <0.001 | 4.58 | 52 | -28 | -6 | Temporal_Mid_R |

| | | | | | | | | |
|---|---|---|---|---|---|---|---|---|
| middle frontal gyrus (MFG) | R | 346 | 0.001 | 4.56 | 42 | 22 | 32 | Frontal_Mid_2_R |
| precuneus | L | 204 | 0.021 | 4.41 | -8 | -66 | 36 | Precuneus_L |
| supplementary motor area (SMA) | R | 196 | 0.025 | 4.17 | 16 | 14 | 66 | Supp_Motor_Area_R |
| **contrast: confCS+ > appCS+** | | | | | | | | |
| insula | L | 869 | <0.001 | 6.37 | -38 | 0 | -10 | Insula_L |
| posterior cingulate cortex (PCC) | R/L | 1515 | <0.001 | 6.23 | -4 | -36 | 22 | Cingulate_Post_L |
| visual association cortex | L | 2022 | <0.001 | 5.92 | -12 | -102 | 4 | Occipital_Mid_L |
| insula | R | 1111 | <0.001 | 5.62 | 40 | 4 | -10 | Insula_R |
| supplementary motor area (SMA) | R/L | 384 | <0.001 | 4.66 | 2 | 20 | 64 | Supp_Motor_Area_R |
| anterior cingulate cortex (ACC) | R/L | 193 | 0.021 | 4.37 | 0 | 16 | 30 | ACC_sup_L |
| **contrast: confCS+ > avCS+** | | | | | | | | |
| lingual gyrus | | 260 | 0.005 | 5.06 | -16 | -90 | 0 | lingual_L |
| lingual gyrus | | 201 | 0.018 | 4.53 | 10 | -86 | -10 | lingual_R |
| **conjunction analysis (confCS+ > neuCS− ∩ avCS+ > neuCS−)** | | | | | | | | |
| middle cingulate cortex (MCC) | R/L | 3062 | <0.001 | 5.67 | 4 | -26 | 28 | Cingulate_Mid_R |
| insula | L | 1031 | <0.001 | 5.32 | -38 | 14 | -6 | Insula_L |
| insula | R | 1356 | <0.001 | 5.19 | 46 | 20 | -8 | Insula_R |
| angular gyrus | L | 559 | <0.001 | 4.91 | -54 | -58 | 34 | Angular_L |
| temporo-parietal junction (TPJ) | R | 784 | <0.001 | 4.9 | 60 | -48 | 30 | Supramarginal_R |
| inferior frontal gyrus (IFG) | R | 270 | 0.007 | 4.87 | 38 | 24 | 28 | Frontal_Inf_Tri_R |
| middle temporal gyrus (MTG) | R | 181 | 0.038 | 4.58 | 64 | -24 | -8 | Temporal_Mid_R |
| pallidum | L | 670 | <0.001 | 4.52 | -8 | 2 | 2 | Pallidum_L |
| precuneus | L | 435 | <0.001 | 4.51 | -8 | -68 | 34 | Precuneus_L |
| calcarine | R | 492 | <0.001 | 4.38 | 14 | -78 | 10 | Calcarine_R |
| superior frontal gyrus (SFG) | R | 188 | 0.033 | 4.12 | 22 | 58 | 20 | Frontal_Sup_2_R |

Note. Only peak coordinates of significant clusters are reported. R/L = bilateral; R = right; L= left

#### Neural activation during response phase (Figure 6)

#### Supplementary Table S9

**Descriptive statistics of family-wise error-small volume correction (FWE-SVC) for local extrema showing greater activity during the response in respective contrast. peak-level $p_{FWE-SVC} < 0.05$. Durations of regressors were set as zero.**

| **Anatomy labeling** | **Hemis-phere** | $p_{FWE-SVC}$ | **Peak T** | **X** | **Y** | Z |
|---|---|---|---|---|---|---|
| **contrast forcedApp > forcedAvd for avCS+** | | | | | | |

| | | | | | | |
|---|---|---|---|---|---|---|
| inferior frontal cortex (IFG) | L | 0.045 | 4.63 | -44 | 14 | 30 |
| middle congulate cortex (MCC) | L | 0.050 | 4.3 | -12 | -32 | 48 |
| insula | R | 0.048 | 4.33 | 38 | -18 | 14 |
| precentral gyrus | L | 0.024 | 4.8 | -44 | 12 | 30 |
| **contrast forcedApp < forcedAvd for neuCS−** | | | | | | |
| insula | R | 0.021 | 4.69 | 40 | 8 | 2 |
| insula | L | 0.046 | 4.38 | -40 | -4 | 0 |
| postcentral gyrus | R | 0.003 | 5.67 | 60 | -20 | 32 |
| postcentral gyrus | L | 0.013 | 5.12 | -52 | -18 | 36 |

Note: Only peak coordinates of significant clusters are reported. R/L = bilateral; R = right; L= left. forcedApp = forced approach; forcedAvd = forced avoidance.

#### Supplementary Table S10

**Descriptive statistics for clusters and local extrema showing greater activity during the response phase in respective contrast at cluster-level $p_{FWE\text{-}corr} < 0.05$ after an initial cluster-defining threshold of $p < 0.001$ across the whole brain.**

| Cluster anatomy (manual labeling) | Hemi-sphere | Cluster size | $p_{FWE\text{-}corr}$ | Peak T | X | Y | Z | Peak label (AAL3) |
|---|---|---|---|---|---|---|---|---|
| **avCS+: forcedApp > focedAvd** | | | | | | | | |
| inferior frontal gyrus (IFG) | L | 309 | <0.001 | 4.74 | -38 | 22 | -12 | Inferior Frontal Orbital Cortex (Region 2) |
| precuneus | R | 250 | 0.003 | 4.34 | 10 | -70 | 38 | Precuneus_R |
| **neuCS−: forcedApp < focedAvd** | | | | | | | | |
| postcentral gyrus | R | 755 | <0.001 | 5.22 | 58 | -18 | 32 | Postcentral gyrus_R |
| cerebelum | R | 197 | 0.006 | 4.93 | 32 | -50 | -26 | Cerebelum_6_R |
| inferior frontal gyrus (IFG) | R | 296 | <0.001 | 4.87 | 46 | -70 | -10 | Temporal_Inf_R |
| middle temporal gyrus (MTG) | L | 312 | <0.001 | 4.77 | -56 | -70 | 6 | Temporal_Mid_L |
| postcentral gyrus | L | 560 | <0.001 | 4.76 | -48 | -20 | 38 | Postcentral gyrus_L |
| insula | R | 292 | <0.001 | 4.73 | 42 | 10 | 2 | Insula_R |
| superior frontal gyrus (SFG) | R | 183 | 0.008 | 4.67 | 26 | -8 | 72 | Frontal_Sup_2_R |

Note: Only peak coordinates of significant clusters are reported. R/L = bilateral; R = right; L= left. forcedApp = forced approach; forcedAvd = forced avoidance.

#### Multivariate pattern analysis (Figure 7E)

Multivariate pattern analysis using meta-analytic ROIs from Neurosynth replicated the conjunction-based results (Table S11), reinforcing the reliability of our findings.

#### Supplementary Table S11

**Results of MVPA-SVR analysis based on ROIs derived from the conjunction analysis of confCS+ > neuCS− ∩ avCS+ > neuCS− and ROIs from Neurosynth.**

| Seed regions (ROIs) | Peak coordinate (MNI) | Sphere size | SVR-correlation | *p* value (Permmutation test with *K* = 5000) |
|---|---|---|---|---|
| **ROIs derived from conjunction analysis of confCS+ > neuCS− ∩ avCS+ > neuCS−** | | | | |
| ACC | [4 36 28] | 5 mm | -0.39 | 0.9446 |
| MCC | [4 -26 30] | 5 mm | -0.07 | 0.4922 |
| **VS** | **[12 10 -6]** | **5 mm** | **0.38** | **0.0228** |
| IFG | [46 22 -10] | 5 mm | 0.02 | 0.3538 |
| insula | [-38 14 -6] | 5 mm | 0.07 | 0.2930 |
| **ROIs derived from Neurosynth meta-analytic maps** | | | | |
| dorsal ACC | [4 32 22] | 5 mm | -0.35 | 0.9148 |
| MCC | [2 -10 42] | 5 mm | -0.48 | 0.9818 |
| **VS** | **[10 10 -6]** | **5 mm** | **0.40** | **0.0216** |
| IFG | [-50 16 12] | 5 mm | -0.002 | 0.3760 |
| anterior insula | [-34 22 0] | 5 mm | -0.23 | 0.7800 |

Note. SVR-correlation means correlation between the predicted values of the SVR model and the true P(avoid) difference between confCS+ and avCS+. Abbreviations: ROIs = regions of interest, MNI = Montreal Neurological Institute, MVPA = multivariate pattern analysis, SVR = support vector regression, ACC = anterior cingulate cortex, MCC = middle cingulate cortex, IFG = inferior frontal gyrus, and VS = ventral striatum. Cluster size for each ROI includes 81 voxels.

#### Functional connectivity during anticipation (Figure 7B)

#### Supplementary Table S12

**Descriptive statistics for clusters and local extrema showing greater functional connectivity during the anticipation in contrast PPI confCS+ > avCS+. Whole-brain cluster-level FWE correction ($p_{FWE\text{-}corr}$ < 0.05) after an initial cluster-defining threshold of $p$ < 0.001 across the whole brain.**

| Seed regions | Peak coordinate (MNI) | Hemisphere | $p_{FWE\text{-}corr}$ | Cluster size | Peak T | X | Y | Z | Cluster anatomy (manual labeling) | AAL3 |
|---|---|---|---|---|---|---|---|---|---|---|
| ACC | [4 36 28] | R/L | 0.003 | 311 | 4.9 | -2 | -20 | -36 | brainstem | Repje_M |
| MCC | [4 -26 30] | — | — | — | — | — | — | — | — | — |
| IFG | [46 22 -10] | R | <0.001 | 802 | 5.82 | 48 | -34 | 34 | temporo-parietal junction (TPJ) | SupraMarginal_R |
| | | L | 0.028 | 193 | 4.77 | -44 | -8 | 22 | temporo-parietal junction (TPJ) | Roladic_Oper_L |

| | | | | | | | | | | |
|---|---|---|---|---|---|---|---|---|---|---|
| | | L | 0.023 | 173 | 6.13 | -24 | -32 | -30 | cerebellum | Cerebellum_4_5_L |
| | | R | <0.001 | 1584 | 5.82 | 54 | -26 | 22 | temporo-parietal junction (TPJ) | Rolandic_Oper_R |
| VS | [12 10 -6] | L | <0.001 | 5177 | 5.72 | -14 | 0 | 56 | superior frontal gyrus (SFG) | Frontal_Sup_2_L |
| | | L | 0.003 | 263 | 4.56 | -24 | -76 | 30 | visual association cortex | Occipital_Mid_L |
| | | L | 0.021 | 177 | 4.49 | -24 | -6 | 22 | caudate | Caudate_L |
| | | R | 0.047 | 145 | 4.12 | 58 | 4 | 14 | temporo-parietal junction (TPJ) | Rolandic_Oper_R |
| | | L | 0.025 | 184 | 5.39 | -24 | -6 | 24 | caudate | Caudate_L |
| insula | [-38 14 -6] | R | <0.001 | 413 | 4.5 | 48 | -36 | 34 | temporo-parietal junction (TPJ) | SupraMarginal_R |
| | | R | 0.016 | 203 | 4.41 | 20 | -40 | 42 | middle cingulate cortex (MCC) | Cingulate_Mid_R |

Note. Only peak coordinates (X, Y, Z) of significant clusters are reported. R/L = bilateral; R = right; L= left. Abbreviations: anterior cingulate cortex (ACC), middle cingulate cortex (MCC), inferior frontal gyrus (IFG), insula, and ventral striatum (VS).
